# Meta-analysis suggests the microbiome responds to Evolve and Resequence experiments in *Drosophila melanogaster*

**DOI:** 10.1101/2020.03.19.999078

**Authors:** Lucas P. Henry, Julien F. Ayroles

## Abstract

**Background:** Experimental evolution has a long history of uncovering fundamental insights into evolutionary processes, but has largely neglected one underappreciated component--the microbiome. As eukaryotic hosts evolve, the microbiome may also evolve. However, the microbial contribution to host evolution remains poorly understood. Here, we re-analyzed genomic data to characterize the metagenomes from ten Evolve and Resequence (E&R) experiment in *Drosophila melanogaster* to determine how the microbiome changed in response to host selection.

**Results:** Bacterial diversity was significantly different in 5/10 studies, primarily in traits associated with metabolism or immunity. Duration of selection did not significantly influence bacterial diversity, highlighting the importance of associations with specific host traits. Additionally, we find that excluding reads from a facultative symbiont, *Wolbachia*, in the analysis of bacterial diversity changes the inference, raising important questions for future E&R experiments in the *D. melanogaster* microbiome.

**Conclusions:** Our genomic re-analysis suggests the microbiome often responds to host selection; thus, the microbiome may contribute to the response of *Drosophila* in E&R experiments. We outline important considerations for incorporating the microbiome into E&R experiments. The E&R approach may provide critical insights into host-microbiome interactions and fundamental insight into the genomic basis of adaptation.

## Background

The microbiome has emerged as a key modulator of many organismal phenotypes [1–3]. While many studies show the impact of the microbiome on host phenotypes, the evolutionary implications remain enigmatic [4–6]. The microbiome may contribute to host evolution in unique ways. First, large effective population sizes and rapid generation times may enable microbes to evolve more rapidly than hosts [7]. Second, the microbiome likely encodes distinct genes compared to the host genome, potentially expanding the genomic reservoir to enable adaptation to diverse selective pressures [3, 8, 9]. If hosts can leverage this microbial evolution, then the microbiome may alter host evolution.

Experimental evolution is a powerful tool to study the basis of adaptation, but remains underutilized in the study of host-microbiome evolution [6, 10, 11]. One particularly well-suited class of these studies is Evolve and Resequence (E&R) experiments [12–14]. E&R experiments build on the long history of using artificial selection in evolutionary biology by incorporating new advances in sequencing technologies to measure the genomic responses to selection. E&R experiments are commonly performed in microbes like *E. coli* or yeast, as well as eukaryotes like *Drosophila* [13]. In general, E&R experiments begin with large outbred populations. The population is reared under a particular selective regime. The selective regime can take many forms, ranging from threshold selection (e.g., egg size) or general survival under some sort of stressor (e.g., low nutrition diets). In parallel, to control for genetic drift, control populations are maintained in a benign (i.e., non-selective) environment. After a number of generations, the control and evolved populations are sequenced to identify regions of the genomes associated with the response to selection. For flies and other eukaryotic hosts, selection is explicitly applied to host populations, but may also act upon the microbiome. When the microbiome influences host phenotypic variation, microbial variation may also affect the response to selection in hosts. Thus, the underappreciated interplay between host and microbial variation has the potential to complicate the interpretation of selection responses based strictly on host genetic variation.

Microbes may be underappreciated drivers of host phenotypic variation. For example, *Wolbachia* infection can rescue deleterious phenotypes in homozygous mutant *Drosophila* lines [15–17]. Body color in aphids is partially determined by *Rickettsia* secondary symbionts [18]. These phenotypic effects are not limited to single microbial species, but also include more complex microbiomes. In cows, the microbiome explained 13% of methane emissions [19] and 26-42% of fatty acid composition of milk [20]. The microbiome also explained 33% of weight gain in pigs [21]. For both pigs and cows, the microbiome contributed almost as much to traits as host genetics. These examples suggest that the microbiome in many host taxa is an important determinant in host phenotypes, and, in turn, may shape the selection response for hosts. E&R experiments may thus be missing a substantial component that shapes the host evolutionary response.

Here, we analyzed the metagenomes from 10 E&R experiments in *Drosophila melanogaster.* Many phenotypes in *D. melanogaster* are responsive to microbial variation, including developmental, metabolic, and immunological traits [22–24]. Furthermore, E&R experiments in *D. melanogaster* capture the evolutionary response to a wide range of different selective pressures, ranging from life history to nutritional to pathogen challenges (Table 1). Thus, E&R experiments in *D. melanogaster* provide a unique opportunity to study how the microbiome responds to host selection. Our goal here is to explore these publicly available data and using meta-analysis, characterize patterns in the metagenomes of these experiments, and to highlighting potential future directions to show how E&Rs can identify signatures of selection in host-microbiome evolution.

**Table 1:** Evolve & Resequence studies analyzed.

| **Pressure** | **Evolved Phenotype** | **Duration of Selection (generations)** | ***Wolbachia* (% reads, min-max)** | **Experimental design (D: diet, S: sequencing)** |
| --- | --- | --- | --- | --- |
| Accelerated development<sup>25</sup> | Flies developed from egg to adult 20% faster than control | 605 | Infected (0.01-1.40%) | D: Banana, corn syrup, agar. S: 25 females (age not reported) pooled from each line; 4 control and 4 evolved. |
| Delayed reproduction<sup>26</sup> | Age of reproduction increased from 28 to 40 days | 50 | Uninfected (0%) | D: Not specified. S: 100 females (age not reported) pooled from each line; 18 control and 18 evolved. |
| Increased lifespan<sup>27</sup> | Median lifespan was increased from 4 weeks to 7-8 weeks | 48 | Uninfected (0%) | D: Agar, yeast, sugar, oatmeal. S: 250 males + 250 females (age not reported) pooled from each line; 3 control and 3 evolved. |
| Egg size<sup>28</sup> | Egg size was selected ~20% larger and smaller eggs | 16 | Infected (0.02-0.11%) | D: Not specified. S: 100 females (age not reported) pooled from each line; 3 control, 3 small, 3 large. |
| Desiccation resistance<sup>29</sup> | Desiccation resistance (hrs until 80% mortality) increased 70-80% | 48 | Infected (70.2-89.7%) | D: Yeast, cornmeal, agar. S: 100 females (age not reported) pooled from each line; 3 control and 3 evolved. |
| Fluctuating temperature<sup>30</sup> | Survival under fluctuating temps 18-28°C daily | 37 | Infected (48.7-75.1%) | D: Standard media. S: 500 females (~7 days old) pooled for each line at different time points; beginning, middle, and end; 3 control and 3 evolved--only compared beginning and end. |
| Salt + cadmium resistance<sup>31</sup> | Survival in constant, spatially, temporally varying salt and/or cadmium | 42 | Infected (0.01-0.03%) | D: Yeast, cornmeal, sugar agar. Diet differed between control and evolved. S: 70 females (age not reported) pooled from each line; 3 control lines and 5 lines for each selection pressure. |
| Starvation resistance<sup>32</sup> | Starvation resistance (hrs to death w/o food) increased ~25% | 83 | Infected (53.0-78.4%) | D: Not specified. S: 100 females (4 days old) pooled for each line; 3 control and 3 evolved lines. |
| Parasitoid resistance<sup>33</sup> | Resistance to parasitoid increased from 20% to 50% | 5 | Uninfected (0%) | D: Not specified. S: 50 females (~5 hrs old) pooled from each line; 16 control and 16 evolved lines. |
| Viral resistance<sup>34</sup> | Resistance to Drosophila C virus increased from 25% to 75% | 20 | Infected (95.1-98.1%) | D: Standard cornmeal-agar. S: 200 individuals (age not reported) pooled from each line; 4 control, 4 procedure control, 4 evolved. |

## Results

The 10 E&R experiments analyzed for metagenomes ranged in a variety of selective pressures (see Table 1 for full description)--from life history (accelerated development [25], delayed reproduction [26], increased lifespan [27], egg size [28]) to abiotic pressures (desiccation resistance [29], fluctuating temperature [30], salt and heavy metal resistance [31]) to biotic pressures (starvation resistance [32], parasitoid resistance [33], viral resistance [34]). All experiments had replicated control and evolved populations, although replication varied from as few as three to as many as 18 (Table 1). Given the importance of the diet in shaping microbial variation, we examined the reported characteristics of the diet from each study. Importantly, for 9/10 studies, the reported diets did not differ between control and evolved populations (Table 1). Only for the cadmium and salt resistance study [31] were diets different for the entire lifespan between control and evolved populations. The starvation resistance study [32] exerted starvation on adults for four days, and then flies that survived were returned to a standard diet to propagate the next generation. While the diets may have varied between studies, only 5/10 studies described the diet (Table 1). The lack of consistent dietary reporting is a major challenge for *Drosophila*-microbiome studies [35]. As the majority of these studies do not report specific dietary information, we are unable to explore the effects of diet across E&Rs in this analysis.

Because each experiment has replicated control and evolved populations, we compared microbiomes within each experiment. In the E&R context, control populations represent the standing genetic variation from which selection proceeds. Thus, by comparing control and evolved populations within experiment, the effects of many different factors (e.g., local laboratory environment, different diets, different fly populations) are controlled for in our analysis. For each experiment, bacterial families were differentially abundant in control and evolved populations (Fig. 1; Supp. Figs 1–10 for individual replicates for each experiment). Control and evolved populations tended to harbor similar bacterial families across replicates, within each experiment, as measured through beta-diversity (Jaccard similarity; Fig. 2, Table 2). Only in two experiments did control and evolved populations differ in community membership-- accelerated development time and delayed reproduction. Bacterial alpha-diversity frequently responded to experimental evolution (Fig. 3). Evolved populations often exhibited reduced levels of bacterial diversity (4/10 studies), though in one case (accelerated development time) bacterial diversity increased (Table 3 for statistical summary). Taken together, the microbiome frequently shifts in response to host selection (i.e., differences in alpha-diversity), but does not necessarily gain different microbes (i.e., no difference in beta-diversity).

**Fig. 1:**
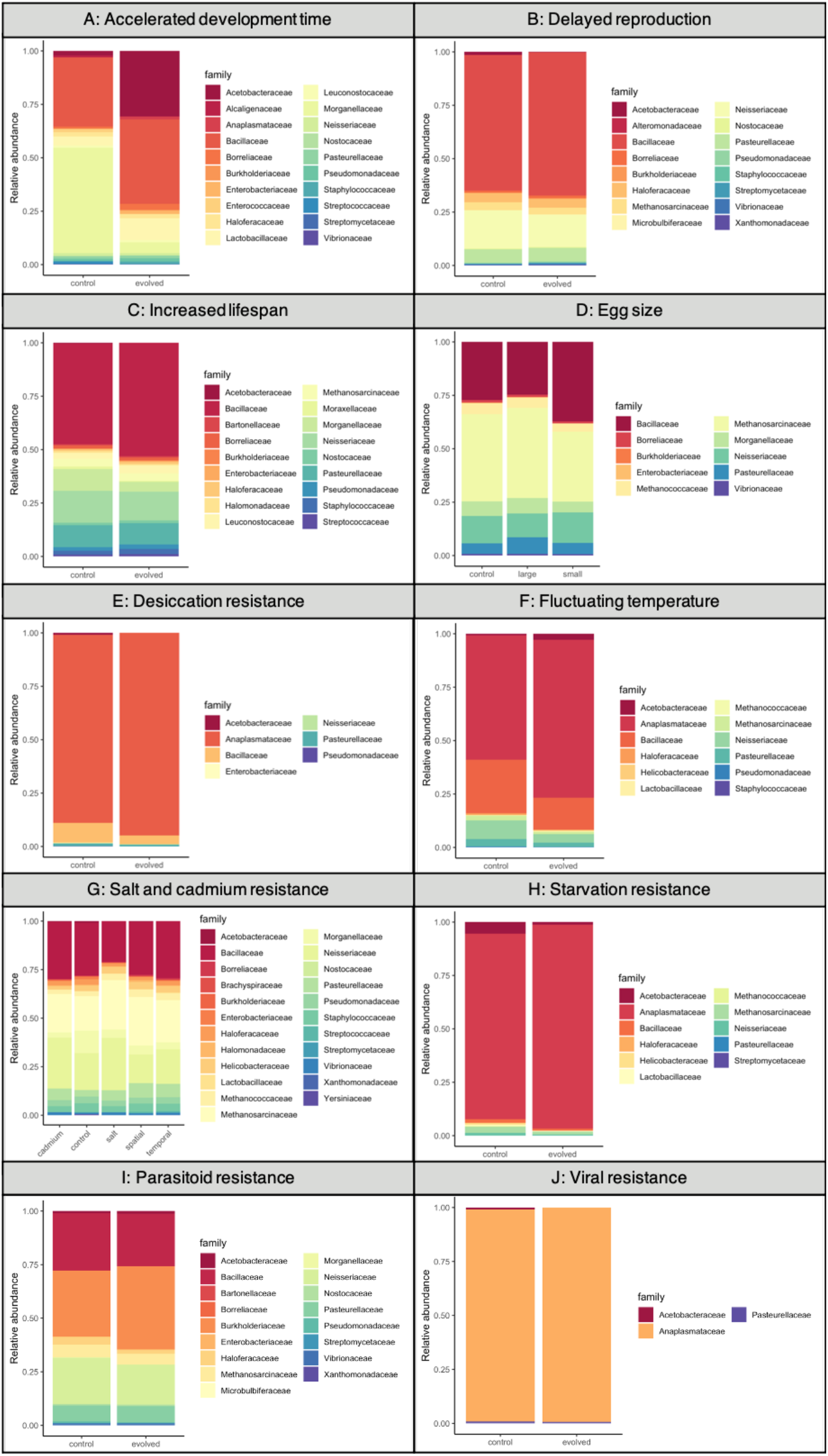
Relative abundance for bacterial families from the 10 E&R experiments. Each experiment was grouped separately; the colors represent different bacterial families in each. Only bacteria that comprised >1% of total reads were visualized.

**Fig. 2:**
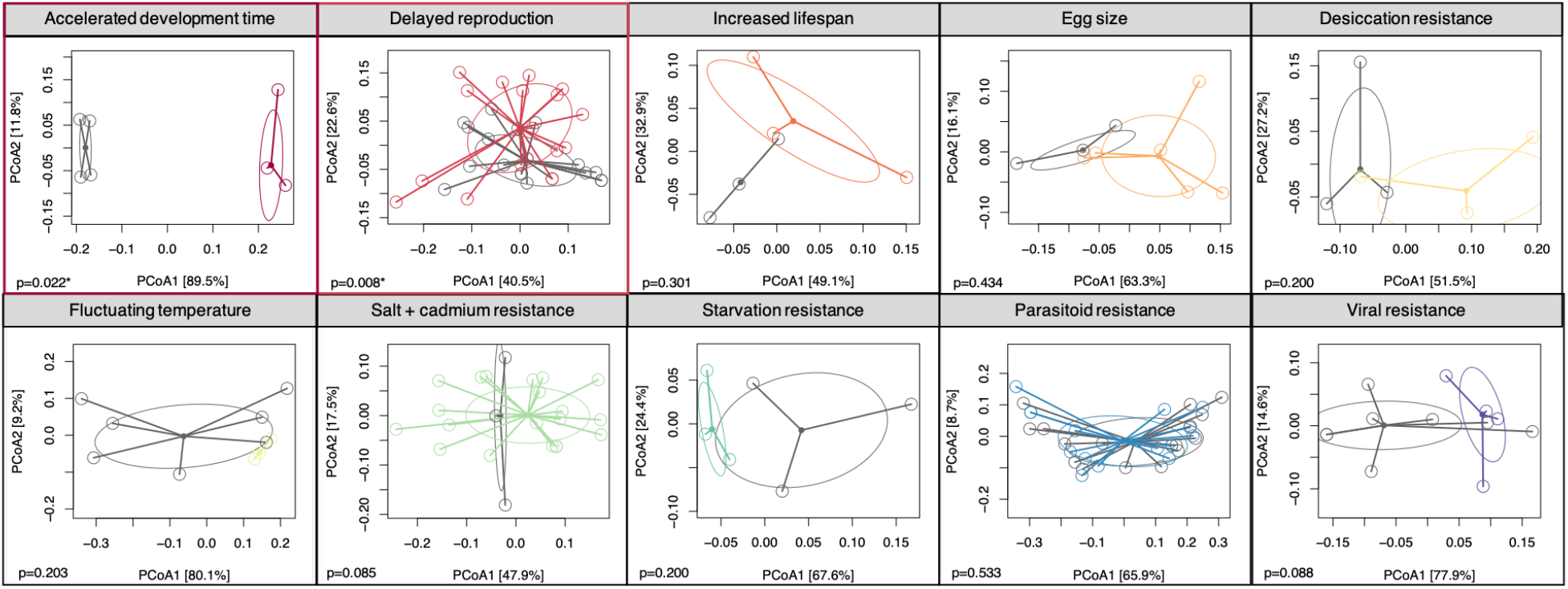
PCoA plots for beta-diversity using Jaccard similarity for the 10 E&R experiments show the majority of control and evolved populations harbor similar bacterial families. Each experiment was grouped separately; grey color represents control populations, and the colored points represent evolved populations. Each sequenced pool is shown as a point with lines connecting to the centroid based on PCoA clustering. The two studies (accelerated development time and delayed reproduction) with significantly divergent microbiomes between control and evolved populations are outlined in colored boxes. P-values are shown for all studies.

**Table 2:**
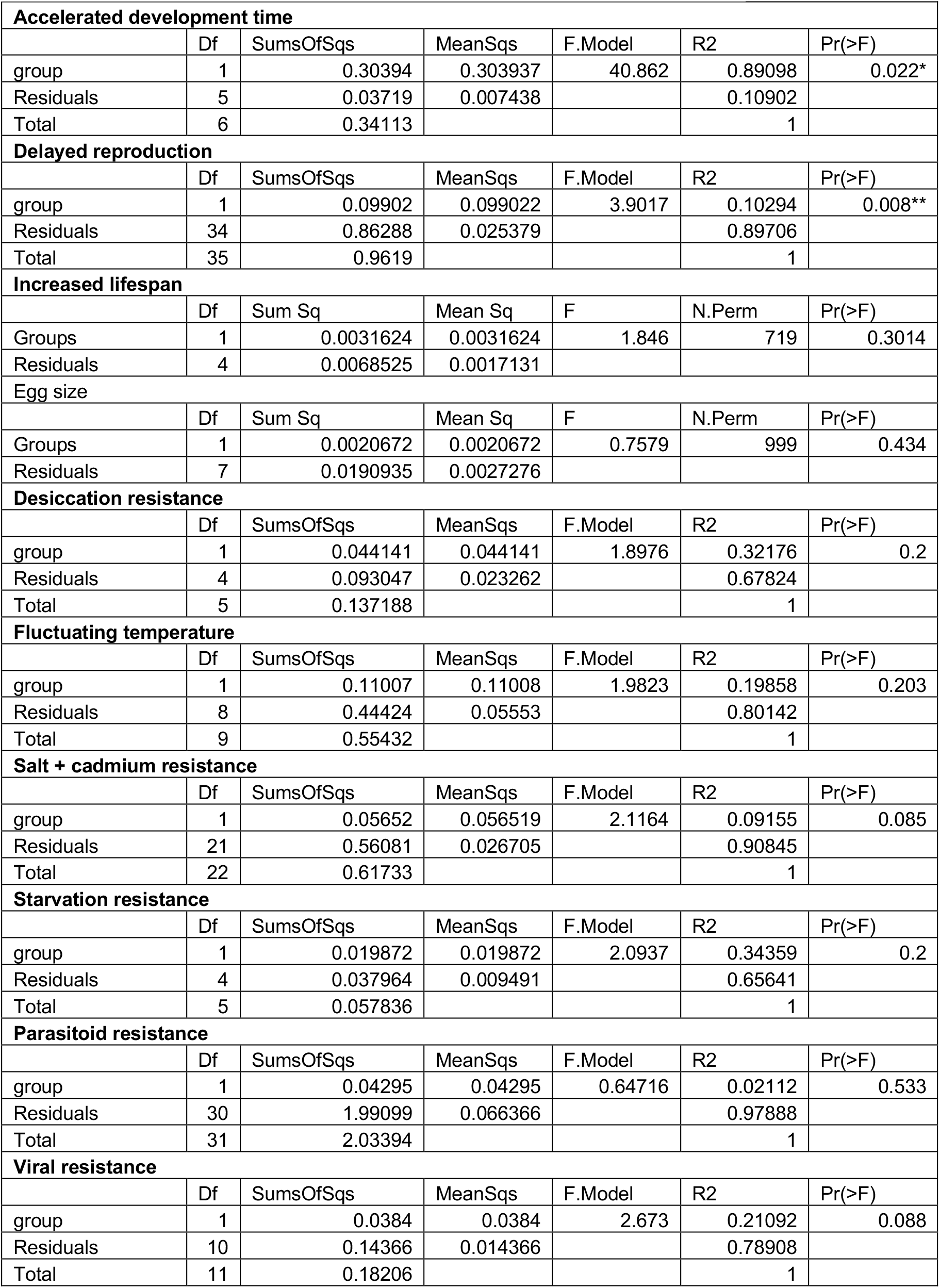
Summary statistics for beta-diversity (Jaccard similarity)

|  |  |  |  |  |  |  |
| --- | --- | --- | --- | --- | --- | --- |
| **Accelerated development time** |  |  |  |  |  |  |
|  | Df | SumsOfSqs | MeanSqs | F.Model | R2 | Pr(>F) |
| group | 1 | 0.30394 | 0.303937 | 40.862 | 0.89098 | 0.022* |
| Residuals | 5 | 0.03719 | 0.007438 |  | 0.10902 |  |
| Total | 6 | 0.34113 |  |  | 1 |  |
| **Delayed reproduction** |  |  |  |  |  |  |
|  | Df | SumsOfSqs | MeanSqs | F.Model | R2 | Pr(>F) |
| group | 1 | 0.09902 | 0.099022 | 3.9017 | 0.10294 | 0.008** |
| Residuals | 34 | 0.86288 | 0.025379 |  | 0.89706 |  |
| Total | 35 | 0.9619 |  |  | 1 |  |
| **Increased lifespan** |  |  |  |  |  |  |
|  | Df | Sum Sq | Mean Sq | F | N.Perm | Pr(>F) |
| Groups | 1 | 0.0031624 | 0.0031624 | 1.846 | 719 | 0.3014 |
| Residuals | 4 | 0.0068525 | 0.0017131 |  |  |  |
| **Egg size** |  |  |  |  |  |  |
|  | Df | Sum Sq | Mean Sq | F | N.Perm | Pr(>F) |
| Groups | 1 | 0.0020672 | 0.0020672 | 0.7579 | 999 | 0.434 |
| Residuals | 7 | 0.0190935 | 0.0027276 |  |  |  |
| **Desiccation resistance** |  |  |  |  |  |  |
|  | Df | SumsOfSqs | MeanSqs | F.Model | R2 | Pr(>F) |
| group | 1 | 0.044141 | 0.044141 | 1.8976 | 0.32176 | 0.2 |
| Residuals | 4 | 0.093047 | 0.023262 |  | 0.67824 |  |
| Total | 5 | 0.137188 |  |  | 1 |  |
| **Fluctuating temperature** |  |  |  |  |  |  |
|  | Df | SumsOfSqs | MeanSqs | F.Model | R2 | Pr(>F) |
| group | 1 | 0.11007 | 0.11008 | 1.9823 | 0.19858 | 0.203 |
| Residuals | 8 | 0.44424 | 0.05553 |  | 0.80142 |  |
| Total | 9 | 0.55432 |  |  | 1 |  |
| **Salt + cadmium resistance** |  |  |  |  |  |  |
|  | Df | SumsOfSqs | MeanSqs | F.Model | R2 | Pr(>F) |
| group | 1 | 0.05652 | 0.056519 | 2.1164 | 0.09155 | 0.085 |
| Residuals | 21 | 0.56081 | 0.026705 |  | 0.90845 |  |
| Total | 22 | 0.61733 |  |  | 1 |  |
| **Starvation resistance** |  |  |  |  |  |  |
|  | Df | SumsOfSqs | MeanSqs | F.Model | R2 | Pr(>F) |
| group | 1 | 0.019872 | 0.019872 | 2.0937 | 0.34359 | 0.2 |
| Residuals | 4 | 0.037964 | 0.009491 |  | 0.65641 |  |
| Total | 5 | 0.057836 |  |  | 1 |  |
| **Parasitoid resistance** |  |  |  |  |  |  |
|  | Df | SumsOfSqs | MeanSqs | F.Model | R2 | Pr(>F) |
| group | 1 | 0.04295 | 0.04295 | 0.64716 | 0.02112 | 0.533 |
| Residuals | 30 | 1.99099 | 0.066366 |  | 0.97888 |  |
| Total | 31 | 2.03394 |  |  | 1 |  |
| **Viral resistance** |  |  |  |  |  |  |
|  | Df | SumsOfSqs | MeanSqs | F.Model | R2 | Pr(>F) |
| group | 1 | 0.0384 | 0.0384 | 2.673 | 0.21092 | 0.088 |
| Residuals | 10 | 0.14366 | 0.014366 |  | 0.78908 |  |
| Total | 11 | 0.18206 |  |  | 1 |  |

**Fig. 3:**
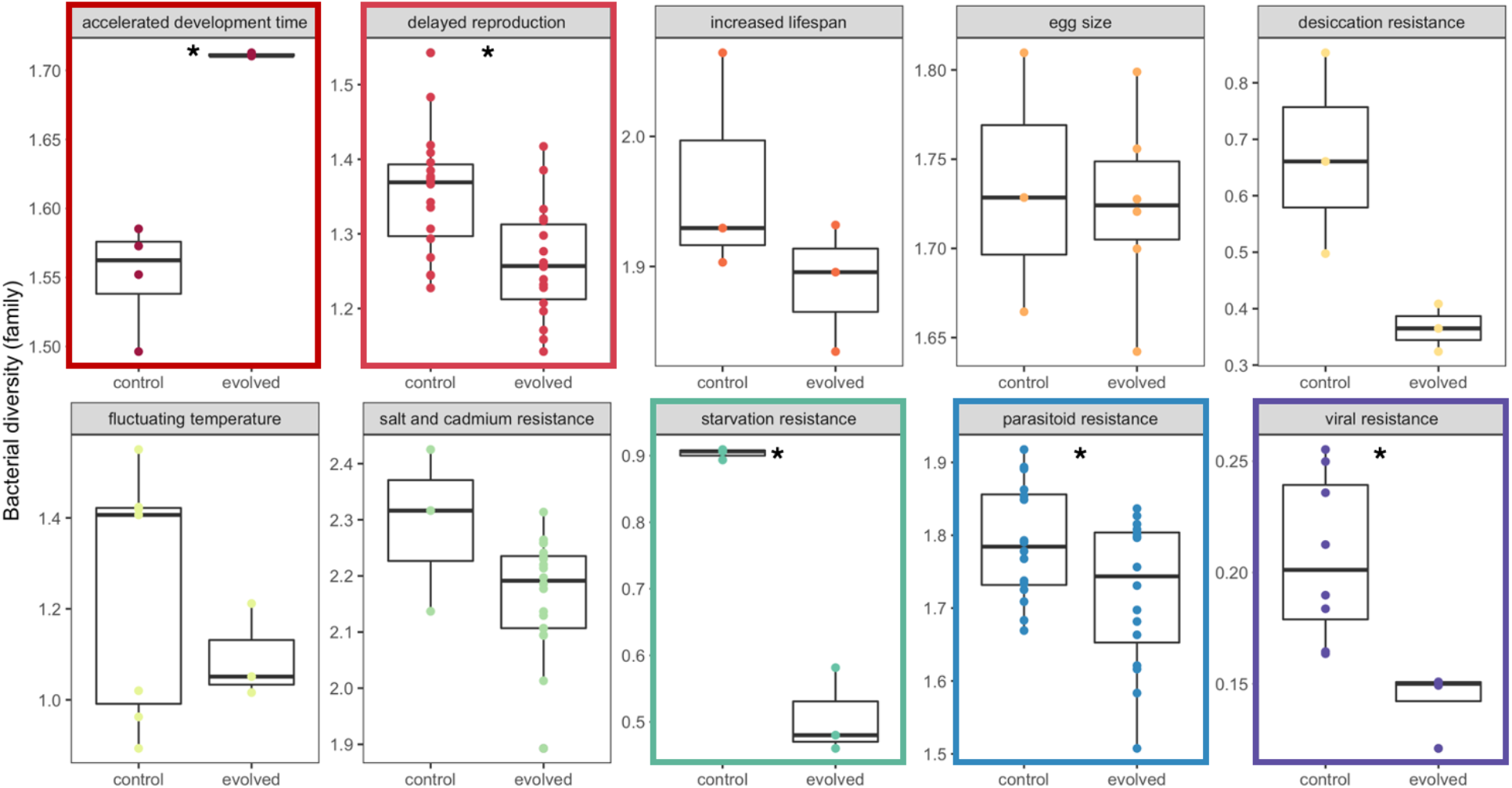
Bacterial diversity between control and evolve populations in 10 E&R experiments. 5/10 experiments had significantly different bacterial diversity (denoted with the colored outline and asterisk). Bacterial diversity was calculated at the family level using Shannon diversity metric. Comparisons between control and evolved populations were within each experiment. Each point represents a pool of sequenced flies showing replication within control and evolved groups, and the details of how many flies/experiments are described in Table 1. Color represents each experiment.

**Table 3:** Statistical differences between control and evolved microbiomes

| Pressure | test stat | df | significance |
| --- | --- | --- | --- |
| accelerated development | t = -8.116 | df = 3.009 | p-value = 0.004 |
| accelerated development without *Wolbachia* | t = -5.511 | df = 3.052 | p-value = 0.011 |
| delayed reproduction | t = 3.558 | df = 33.652 | p-value = 0.001 |
| increased lifespan | t = 1.364 | df = 3.172 | p-value = 0.261 |
| egg size | t = 0.213 | df = 3.105 | p-value = 0.845 |
| egg size without *Wolbachia* | t = 0.250 | df = 3.077 | p-value = 0.818 |
| desiccation resistance | t = 2.883 | df = 2.224 | p-value = 0.090 |
| desiccation resistance without *Wolbachia* | t = 0.394 | df = 3.985 | p-value = 0.714 |
| fluctuating temperature | t = 1.238 | df = 7.996 | p-value = 0.251 |
| fluctuating temperature without *Wolbachia* | t = -4.135 | df = 6.886 | p-value = 0.005 |
| salt and cadmium resistance | t = 1.431 | df = 2.287 | p-value = 0.274 |
| salt and cadmium resistance without *Wolbachia* | t = 1.431 | df = 2.287 | p-value = 0.274 |
| starvation resistance | t = 10.448 | df = 2.065 | p-value = 0.008 |
| starvation resistance without *Wolbachia* | t = -1.413 | df = 2.903 | p-value = 0.255 |
| parasitoid resistance | t = 2.179 | df = 28.394 | p-value = 0.038 |
| viral resistance | t = 4.265 | df = 9.829 | p-value = 0.002 |
| viral resistance without *Wolbachia* | t = -2.851 | df = 7.193 | p-value = 0.024 |

Because the number of generations varied across E&R experiments (from 5-605 *Drosophila* generations; see Table 1), we also tested if change in microbial diversity was correlated with duration of host selection. One hypothesis is that shorter selection experiments provide less opportunity for the microbiome to change, while longer selection provides more opportunity for increased microbial change. The change in microbial diversity was not correlated with duration of selection after controlling for each study as a random effect (Fig. 4, r=0.05, p=0.649). The specific nature of the selective pressure appears to be more important in driving changes in the evolving microbiome as experiment explained 76% of variance in our model (Table 4). For example, the evolved microbiome in the starvation resistance experiment exhibited the greatest change in bacterial diversity. This may not be surprising given that the *Drosophila* microbiome has been shown to be tightly linked to the regulation of metabolic networks [22]. For other traits, like egg size, the microbiome did not significantly respond to experimental evolution. This analysis suggests that the effect of selection on microbiome is likely trait specific.

**Fig. 4:**
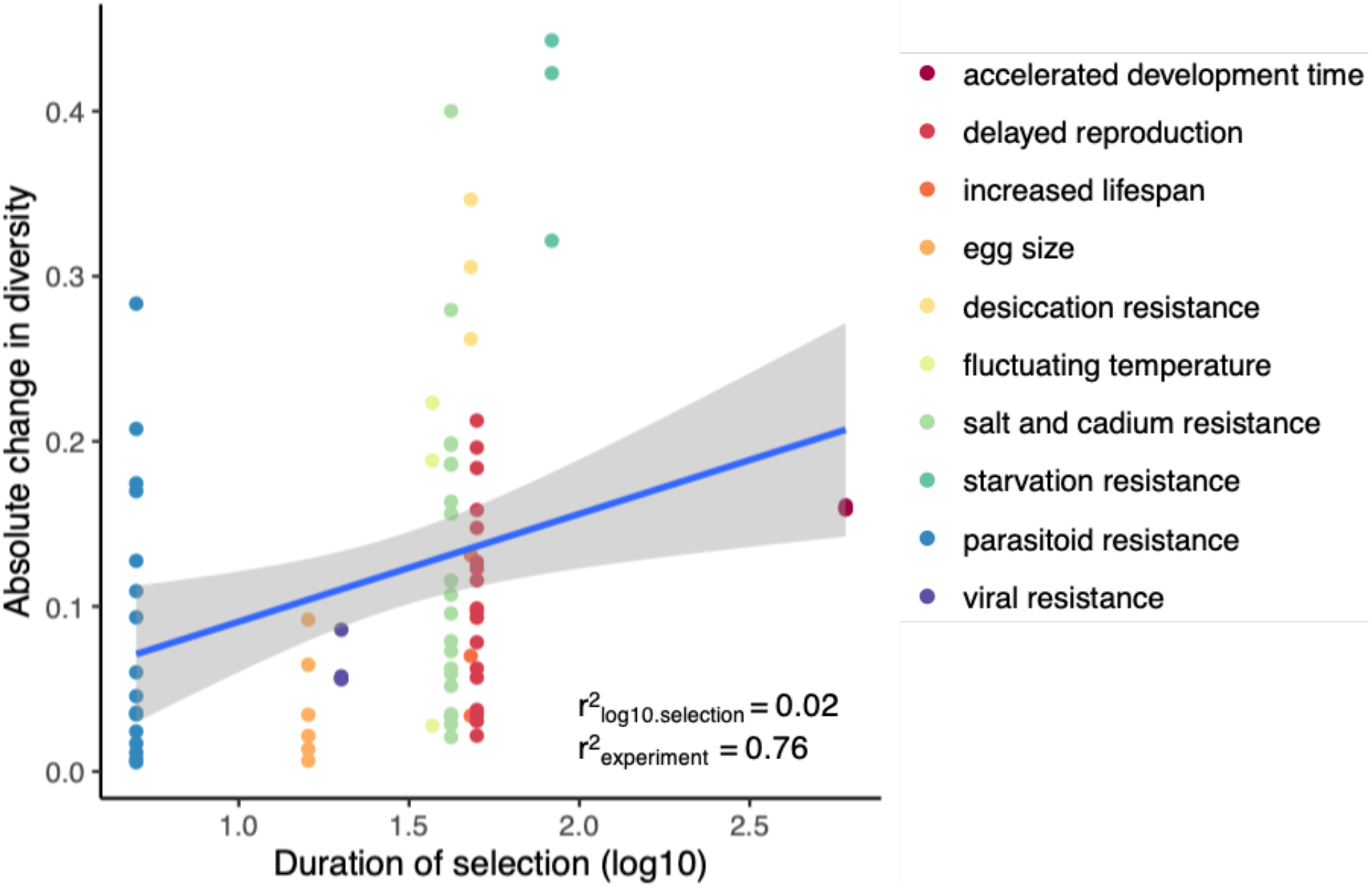
Evolved bacterial diversity was not significantly correlated with duration of selection after controlling for differences between experiments as random effects (*r* = 0.05, p=0.649). Random effects (experiment) explained 76% of variation (see Supp. Fig. 11). Generations of selection range from 5 generations (parasitoid resistance) to > 500 generations (accelerated development). Each point represents the difference between average control diversity and each pool of evolved flies for each experiment. Points are colored by experiment.

**Table 4:** Summary statistics of mixed model to assess the relationship between duration of selection and change in diversity. Random effect was modeled as experiment (i.e., accelerated development time, starvation resistance, etc.).

| Fixed effects |  |  |  |
| --- | --- | --- | --- |
|  | Estimate (std. error) | T-value | Pr (> \| t \|) |
| Intercept | -0.185 (0.162) | -1.137 | 0.291 |
| log10.length | 0.046 (0.097) | 0.473 | 0.649 |
| Random effects |  |  |  |
|  | Variance | Std. deviation |  |
| Experiment | 0.0225 | 0.1500 |  |
| Residual | 0.007 | 0.084 |  |
| Summary |  |  |  |
| Observations |  | 79 |  |
| Log Likelihood |  | 66.128 |  |
| Akaike Inf. Crit. |  | -124.256 |  |
| Bayesian Inf. Crit. |  | -114.779 |  |

We note that, in seven out of ten studies, flies were infected with *Wolbachia* (Table 1). *Wolbachia* was <2% of reads for three studies, but 48-98% for the other four studies. To better understand the association between *Wolbachia* and the microbial response to selection, we focused on these four studies with high relative abundance (Fig. 5). *Wolbachia* was significantly more abundant for evolved populations in starvation and viral resistance, though also tended to increase for desiccation resistance and fluctuating temperatures (Table 5 for statistical summary). Furthermore, excluding *Wolbachia* reads from our analysis revealed different inferences about the response of the evolved microbiome (Fig. 6, Table 3 for statistical summary). First, in desiccation resistance, excluding *Wolbachia* reads reduced the difference in diversity between control and evolved populations, but did not change our inference as no significant difference was observed for the full bacterial community. Second, excluding *Wolbachia* reads leads to an increase in environmentally acquired bacterial diversity in the evolved lines under fluctuating temperatures. Third, excluding *Wolbachia* reads shows that environmentally acquired bacteria did not respond to selection in starvation resistance as there was no difference between control and evolved lines. Finally, in viral resistance, excluding *Wolbachia* increased environmentally acquired bacterial diversity in evolved lines, while the whole community showed reduced diversity. For the three studies with low *Wolbachia* abundance, excluding *Wolbachia* reads did not impact estimates of diversity (Table 3). Taken together, importantly, we suggest careful consideration towards considering inclusion or exclusion of *Wolbachia* reads, as it will significantly alter the observed response to selection in the microbiome.

**Fig. 5:**
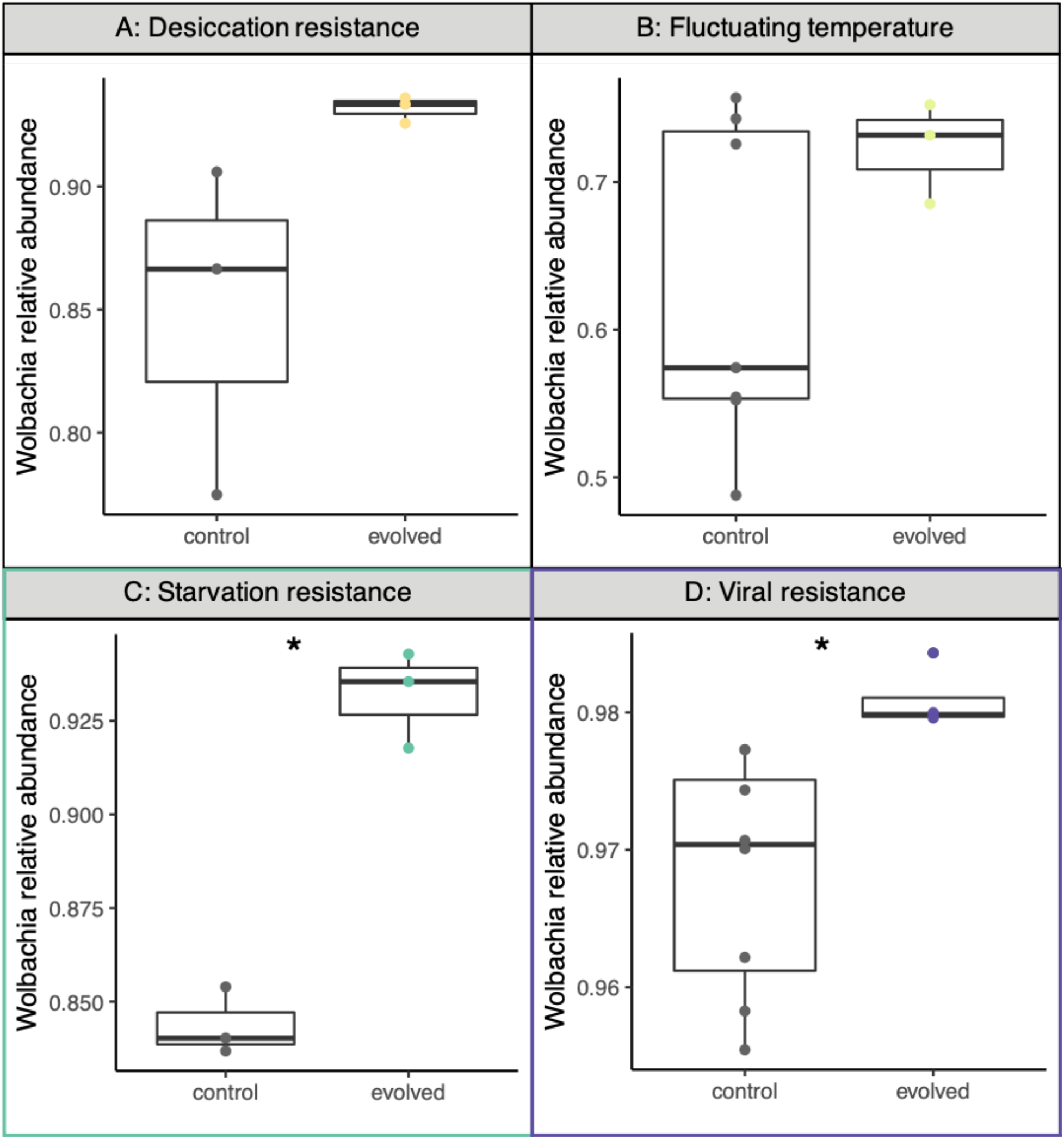
*Wolbachia* relative abundance tended to increase in evolved populations for four studies, but was only statistically significant in starvation resistance and viral resistance (denoted with colored box and asterisk). Comparisons were made within each study between control and evolved populations. Each point represents a pool of flies.

**Table 5:** Statistical differences in *Wolbachia* relative abundance between control (C) and evolved (E) populations.

| Experiment | C min | C max | C avg | E min | E max | E avg |
| --- | --- | --- | --- | --- | --- | --- |
| **Desiccation resistance** |  |  |  |  |  |  |
| Relative abundance | 0.775 | 0.906 | 0.849 | 0.926 | 0.936 | 0.932 |
| Kruskal-Wallis $X^2 = 3.8571$, df = 1, p-value = 0.04953 | | | | | | |
| **Fluctuating temperature** |  |  |  |  |  |  |
| Relative abundance | 0.488 | 0.757 | 0.628 | 0.685 | 0.752 | 0.723 |
| Kruskal-Wallis $X^2 = 1.0519$, df = 1, p-value = 0.3051 | | | | | | |
| **Starvation resistance** |  |  |  |  |  |  |
| Relative abundance | 0.837 | 0.854 | 0.844 | 0.918 | 0.943 | 0.932 |
| Kruskal-Wallis $X^2 = 3.8571$, df = 1, p-value = 0.04953 | | | | | | |
| **Viral resistance** |  |  |  |  |  |  |
| Relative abundance | 0.955 | 0.977 | 0.968 | 0.98 | 0.984 | 0.981 |
| Kruskal-Wallis $X^2 = 7.3846$, df = 1, p-value = 0.006578 | | | | | | |

**Fig. 6:**
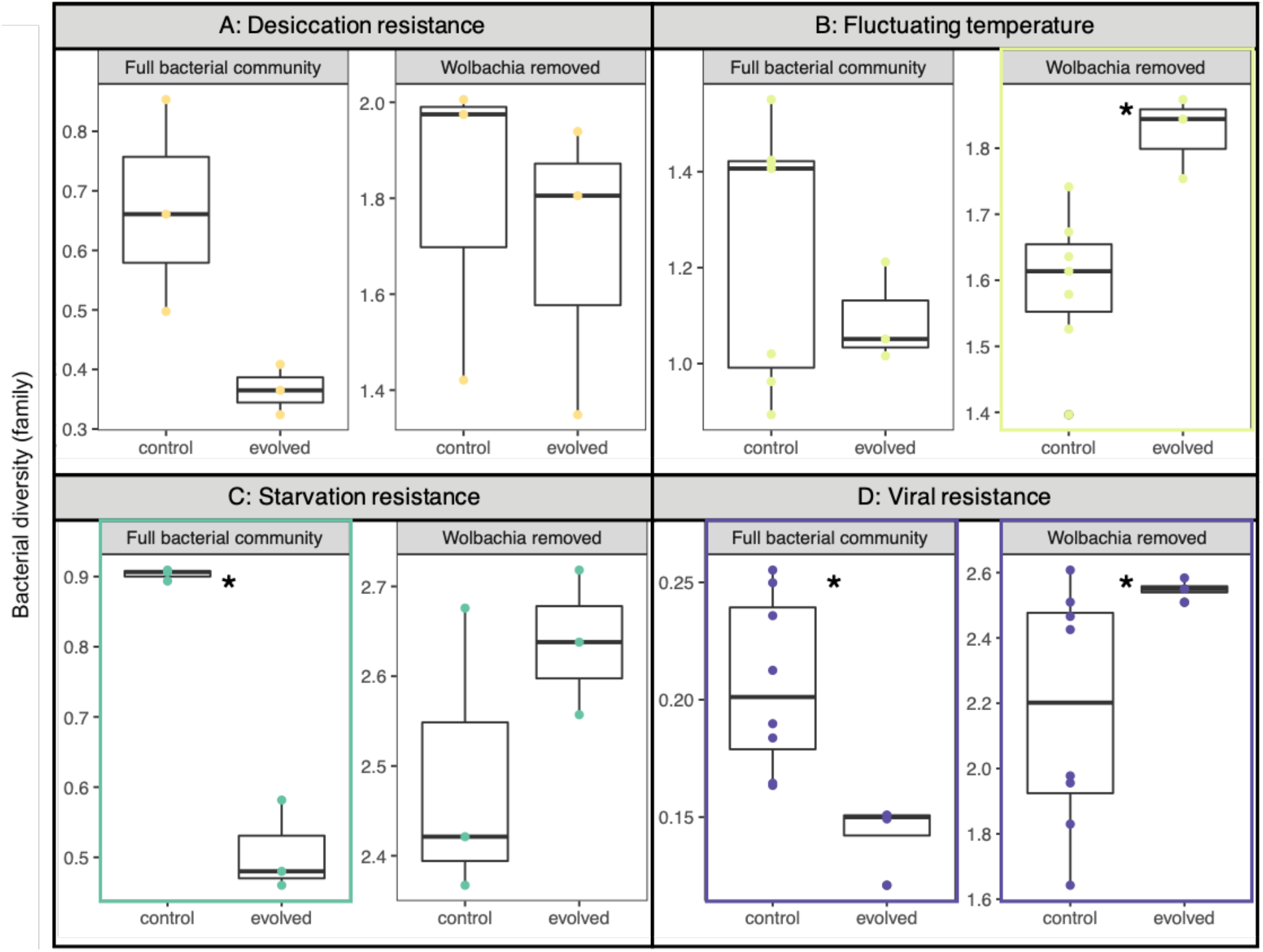
Differences in bacterial diversity when including (full bacterial community) or excluding *Wolbachia* reads in four of the E&R experiments analyzed. Each point represents bacterial diversity for a pool of sequenced flies. Scale bars are different because diversity comparisons are within either the full community or *Wolbachia* removed groupings for each experiment. The four experiments highlight how the difference between control and evolved microbiomes depends on whether *Wolbachia* is included. For example, for starvation resistance, diversity for the whole community is reduced when *Wolbachia* is included. However, when *Wolbachia* is excluded, there is no response in bacterial diversity between control and evolved populations. Full summary of differences can be found in Table 2.

## Discussion

To our knowledge, this is the first systematic examination of the microbiome in E&R experiments in *D. melanogaster*. Given the many fundamental insights gained from *Drosophila* in E&R experiments [13], our results here uncover another layer of variation previously unexplored--the microbiome. The microbiome changed under some selective pressures, while it was unaffected by others (Fig. 1, 3). Pressures closely linked to metabolic processes, like starvation resistance or development time, or immunity affected microbial diversity the most. In *Drosophila*, bacterial genes that increase glucose assimilation and fat storage are necessary for bacterial establishment in the host gut, suggesting that hosts select for bacteria to enhance metabolism [36–38]. Other pressures, like selection for increased lifespan, egg size, or abiotic stressors (e.g temperature and heavy metals), did not substantially impact microbial diversity (Fig. 3). It is not surprising that not all selection pressures shape the microbiome; indeed, in *Drosophila*, traits such as activity level, sleep, and some aspects of nutrition are known to not be influenced by the microbiome [39–42]. Our results suggest that the microbiome changes along with host evolution in the E&R context – although we emphasize that the data we present are a re-analysis of existing genomic data and not derived from new manipulative experiments. Our results here contribute to a growing body of literature suggesting that when the microbiome contributes to host phenotypic variation, the microbiome has the potential impact host evolutionary trajectories [6, 43]. The evolutionary interplay between host and their microbiome may play an important role in driving host evolution (beyond the explicit selective pressures exerted in these experiments), and this level of variation should not be ignored in E&R analyses.

We observed several generalities in the microbial response in E&R experiments. First, the microbiome in both control and evolved populations was composed of similar bacterial families (Fig. 1), suggesting selection did not lead to the complete replacement by different bacterial taxa in evolved populations. In evolved populations, only a few of the bacterial families increased in relative abundance. Furthermore, in all studies, replicate lines from both control and evolved populations show similar community compositions, suggesting consistent effects on the microbiome (Figure 2, Supp. Figs. 1–10). Bacteria that contribute to the host adaptation may be more likely to persist under the selective pressure, increasing in abundance and facilitating local adaptation. Second, the increase in abundance of particular bacterial families also contributed to the frequent reduction in diversity. The reduction in diversity likely reflects local adaptation in the microbiome, but potentially also the loss of genetic diversity in the host. We expect that the rapid nature of E&R experiments, combined with strong selective pressures, results in lower heterozygosity levels across the genome following selection in E&R experiments [13]. Host genetics shapes a significant fraction of the fly microbiome [44], and perhaps the loss of diversity in the host genome also contributed to the reduction in microbial diversity observed here. While evolved populations have reduced heterozygosity, they still maintain substantial heterozygosity across the genome. More research is necessary to understand how host genome-wide diversity affects microbial diversity, or if only certain host loci are the key drivers of changes in microbial diversity. Uncovering the specific host genetic loci that may be associated with microbial changes is beyond the scope of this current study; however this is an important factor to consider in future studies.

For the evolved microbiomes, bacteria may have evolved different functions that hosts can leverage. For example, the relative abundance of *Acetobacteraceae* is enriched in the evolved populations for accelerated development time (Fig. 1A). *Acetobacter* produces acetic acid that modulates the insulin/insulin-like signalling (IIS) growth factor pathway in flies [36]. More so, *Acetobacter* is frequently associated with accelerated development compared to other bacteria [24, 36, 45, 46]. The IIS pathway may also integrate metabolic products from other bacteria in the microbiome to help regulate fly metabolism. *Wolbachia* infection has been shown to increase insulin signaling in *Drosophila* [17]. The increased *Anaplasmataceae* abundance in the evolved populations may better regulate metabolic traits to mitigate selection in the starvation resistance experiment (Fig. 1H). We hypothesize that increased relative abundance for particular bacteria in the evolved populations corresponds to functional changes and is suggestive of fitness benefits for the fly. Subsequently, flies transmit and preferentially associate with the beneficial microbes. However, bacteria may also be increasing in the evolved conditions independently of any host selection. To better understand how microbial evolution interacts with host evolution, longitudinal sampling over the course of the evolutionary trajectory is necessary. Identifying if beneficial adaptations emerge first in the microbiome and then alter allele frequencies in host populations would provide key insights into how host-microbiome interactions shape eukaryotic evolutionary processes.

The temporal aspect of host-microbiome evolution is important, but underexplored and thus poorly understood. Our analysis suggests that time did not significantly affect the difference in diversity between control and evolved populations (Fig. 4). This might be because the microbiome changes rapidly, within a single host generation, but the evidence for rapid change is inconsistent. One study found that the microbiome was significantly different when flies were starved or shifted to a high fat diet [47], while another study found no differences when shifted to low or high sugar diets [48] within their lifespan. Finally, in flies mono-associated with *Lactobacillus* reared in nutrient poor diets, *Lactobacillus* evolved beneficial mutations that promoted fly growth in only 5 fly generations [49]. While most experiments did not manipulate diets in our E&R analysis, the findings from these three studies suggest a range of timescales in which the microbiome may evolve. Conducting longitudinal surveys of the microbiome during experimental evolution is essential to understanding if and how the microbiome shapes host evolutionary trajectories.

Excluding *Wolbachia* reads from the analysis had substantial effects on the inference for the observed response in the microbiome (Fig. 6). This change in inference often resulted from an increase in *Wolbachia* abundance in evolved populations (Fig. 5). The phenotypic effects exerted by *Wolbachia* on their hosts often depend on the degree of increase in abundance. For example, higher abundance provided stronger cytoplasmic incompatibilities [50], increased protection from viruses [51], or greater reductions in lifespan [52]. However, several factors may actually confound the *Wolbachia* results observed in our meta-analysis here. Infection was only assessed from pools of flies – we do not have access to individual level status (Table 1). Relative abundance may reflect the average relative abundance within individuals or heterogeneous infection patterns across individuals. We believe this second scenario of heterogeneous infection across individuals is not likely for the four high *Wolbachia* studies we examined in more detail. In a study that examined how *Wolbachia* infection spreads within outbred populations, *Wolbachia* infection across individuals increased to 80% by 10 generations and nearly 100% by 32 generations [53]. This suggests that for the four high Wolbachia studies, most individuals were infected. However, for the three studies with low *Wolbachia* abundance, it may be that *Wolbachia* infection status is highly heterogeneous across individuals or reflects recent *Wolbachia* infections (or simply contamination). Fly age also affects *Wolbachia* abundance, increasing in older flies [54]. Unfortunately, age of flies was only described in 3/10 studies (Table 1), but was always similar between control and evolved populations. Age might affect our results if control and evolved populations were systematically collected at different ages. We think this unlikely as it would also bias the genomic analysis as survival may differ between control and evolved populations at different ages (e.g., viability selection). Overall, designing experiments that explicitly control for *Wolbachia* infection is necessary to understand its potential influence on host-microbiome evolution [14, 23].

Importantly, the data presented here cannot answer the question whether *Wolbachia* influences the microbiome during fly evolution--rather, our analysis highlights several complications of *Wolbachia*. There are both practical and biological reasons to exclude *Wolbachia* from most microbiome studies as it is intracellular, low abundance in the gut, has complex effects on host traits, overrepresentation in 16S rRNA profiling--all characteristics distinct from the more common bacteria in the fly microbiome, like *Acetobacter* and *Lactobacillus* [23, 55, 56]. Indeed, many studies in *D. melanogaster* either only use uninfected flies [23, 39, 45, 46] or computationally remove *Wolbachia* reads during 16S rRNA microbiome analysis [24, 57–60]. We emphasize the importance of considering and testing the potential influence of *Wolbachia* on fly adaptation and trait variation. In the studies analyzed here, 7/10 studies used flies infected with *Wolbachia*, but only two explicitly mention *Wolbachia* infection status in their flies--fluctuating temperature [30] and viral resistance [34]. Furthermore, only Martins et al. [34] estimated the effects of Wolbachia on host phenotype by clearing *Wolbachia* infections and then re-measuring phenotypes. Notably, no study assessed the change in *Wolbachia* relative abundance. While our approach cannot assess the functional importance, future studies could take a similar approach to Martins et al. by clearing *Wolbachia* infections and re-phenotyping. We acknowledge there is currently ambiguity about including *Wolbachia* in the microbiome as discussed here. We raise these points to highlight unintended potential complications of *Wolbachia* infection, particularly for incorporating the microbiome into E&R studies.

Furthermore, *Wolbachia* has a variety of effects on fly biology, ranging from reproductive phenotypes to immunity to nutrition [61–63] and may substantially influence *Drosophila* evolution [61, 64]. *Wolbachia* may also affect the microbiome. In a comparison of a single genotype of flies infected and uninfected with *Wolbachia*, uninfected flies had twice as much *Acetobacter* [55]. However, in the same study, a different fly genotype did not display this effect. Yet, another study found that *Wolbachia* infection increased *Acetobacter* abundance [56]. These effects are inconsistent and likely depend on interactions between fly genotype, *Wolbachia* genotype, and environmental conditions. If *Wolbachia* interacts positively or negatively with different bacteria, then *Wolbachia* may also influence how the microbiome shapes host phenotypes and contributes to the host evolutionary trajectory. Taken together, the interactions between *Wolbachia*, host, and microbiome are likely complicated. Many insects are infected with *Wolbachia* or similar intracellular symbionts, and these microbe-microbe interactions may have important implications for the host [65].

While this is the first examination of the microbiome in the E&R context, other studies have implicated the microbiome in host adaptation in *D. melanogaster*. For example, as previously mentioned, when flies were monoassociated with *Lactobacillus plantarum* in nutrient poor environments, *L. plantarum* rapidly evolved symbiotic benefits to increase fly fitness [49]. Across replicates, the *de novo* appearance of several SNPs in the acetate kinase gene (*ackA*) in *L. plantarum* promoted larval growth and nutrition, and subsequently, this *L. plantarum* variant increased in frequency across fly generations. In another study, microbiome manipulation shifted allele frequency in seasonally evolving *D. melanogaster* to match latitudinal patterns of fly genetics [59]. Both of these studies rely on mono-associations with single microbes, but this likely does not realistically capture host-microbe dynamics. Higher order interactions among bacteria shape phenotypes in *Drosophila* [45, 46]. Interaction among microbes, like cross-feeding of metabolites between *Acetobacter* and *Lactobacillus*, can enable mutually beneficial growth for both bacteria species as well as increases bacterial growth, but critically also alters fly metabolism [66]. Furthermore, even strain-level variation within a bacterial species can have divergent effects on host phenotypes [67, 68]. The technical challenges associated with accurately quantifying genetic variation across complex microbial populations necessitated these mono-association experiments. Fortunately, new emerging methods are enabling the identification of signatures of selection in complex microbiomes [69, 70]. Future experiments with more complex and realistic microbiomes will show how microbe-microbe interactions contribute to host adaptation.

Taken together with our analyses, both the host and microbiome likely evolve in response to selection. More generally, other systems like *Brassica* and *Arabidopsis* have also shown that selection on hosts changes the microbiome as well [71, 72]. In both these studies, transplanting an evolved microbiome into unevolved hosts changed host phenotypes, suggesting that the microbiome has the capacity to transfer adaptive potential. Similar approaches could be applied to *Drosophila* following E&R experiments. Combined with the rich genetic resources and experimental ease in *Drosophila*, microbiome transplants could illuminate key processes underlying host-microbiome evolution.

We note the experiments analyzed here were not designed explicitly to test the role of the microbiome in host adaptation. This may impact our results in several ways. None of these studies were executed with quality control measures that can affect estimates of microbial diversity, such as process blanks during DNA extraction, no template controls during PCR, and batch effects during library preparation [69, 73–75]. While we applied an arbitrary cutoff to remove contaminants, it is difficult to know how potential contaminants may affect the observed results. However, contamination would have to differentially affect control and evolved microbiomes to influence our results--which we believe is unlikely. Surveys of microbial diversity in *D. melanogaster* typically use 16S rRNA profiling and find bacteria from the *Acetobacteraceae*, *Firmicutes*, and *Enterobacteriaceae* [22, 24, 76]. Our mapping approach detected these bacteria commonly associated with *D. melanogaster*, but also found abundant methanogens and human commensal microbes (Fig. 1). One discrepancy could arise from our metagenomic approach, which will often lead to different conclusions than 16S rRNA profiling [69]. Mining metagenomes from existing whole genome sequencing is an emerging area of research in the microbiome, and more work is necessary for biological interpretations [5, 69]. Finally, none of the flies sequenced in these studies were surface sterilized, and thus, the metagenomes characterized here result from both the external body surface and internal gut microbiome. However, the external microbiome is orders of magnitude less abundant than the internal microbiome across the fly lifespan [77]. While we cannot distinguish between external and internal microbiomes in this analysis, future studies should be clear if the total (external and internal) or gut microbiomes were sequenced. Nevertheless, the consistent differences in the microbiome across experiments shown here highlight how E&R experiments could provide exemplary opportunities to investigate the genetic basis underlying host-microbiome evolution.

## Conclusions

For researchers interested in adapting the E&R approach for host-microbiome interactions, we have several key recommendations. As we have suggested above, more intensive temporal sampling to capture both microbial and host evolution is necessary. For *Drosophila*-microbiome E&R experiments, researchers may wish to begin the experimental evolution by standardizing the microbiome between control and evolved populations, like with the 5-species bacterial community commonly used [23, 78] or fly feces to mimic natural, but standardized microbial inoculation [79, 80]. Second, as much of the microbiome is determined by the environment in flies, researchers need to use consistent brands of yeast, preservatives, and other aspects of diet/environment. For example, different preservatives have different effects on the microbiome and behavioral traits [40, 41]. *Drosophila* in different labs in the same building (with the same kitchen for fly food) had different microbiomes [76], suggesting that several aspects of the environment are important in shaping the fly microbiome. Finally, as outlined by Goodrich et al. [81], microbiome research requires careful planning (with both biological and technical controls), extensive documentation, and consistency. Importantly, we are not advocating that every E&R experiment incorporates the microbiome, but note that the microbiome may impact conclusions from E&R experiments. For researchers not explicitly interested in the microbiome, our primary recommendation is to clear fly lines of *Wolbachia* to avoid potential confounding effects between host genetic and *Wolbachia* evolution as others have suggested [14, 23].

In conclusion, the microbiome frequently responded to selection in ten E&R studies in *D. melanogaster.* Our results here associate the microbiome in the host response to some selective pressures, but more work is necessary to partition the relative effects of host genetics and microbial evolution. We observed large differences in bacterial diversity between control and evolved populations, but a key question remains--if and how the microbiome alters the host response to selection. Combining E&R experiments with approaches from quantitative genetics will be especially fruitful to dissecting the microbial contribution to host evolution [6]. Tracking the rate of microbial evolution over multiple timepoints during fly adaptation will be particularly helpful to elucidate whether the microbiome shapes the host evolutionary trajectory. Partitioning the microbial effects on host phenotype during adaptation may show that microbiome facilitates or impedes host adaptation. Reciprocal transplants over the course of host adaptation will also demonstrate how the microbiome modifies host evolution. Our results here suggest that the microbiome might influence host evolution, but do not prove it. To measure how the microbiome affects host evolutionary trajectory, combining several of these techniques will be necessary. Overall, incorporating the microbiome into E&R experiments will provide fundamental insights into host-microbiome evolution.

## Methods

We searched the literature for E&R experiments in *D. melanogaster* where replicated control and selection lines were derived from outbred populations and raw. fastq data were publicly available. We found 10 studies that met these criteria. Our analyses captured a wide range of different selection pressures, from life-history traits to abiotic and pathogen pressures, enabling generalizations about the microbiome response to host selection. In all cases, the E&R approach sequenced pools of individuals from different selection regimes, but each E&R study had different levels of replication (summarized in Table 1). We report the diet as described in the publication for each study (Table 1). While most studies did not publish specifics about the diet, we noted the diets that differed between control and evolved populations (only one study [31]); if the publication did not specify, we reasonably assumed diets were the same. These were the only data available from published E&R experiments in *D. melanogaster* at the time of submission.

Raw sequences were cleaned using Trimmomatic [82] to remove sequencing adapters, remove low quality reads (average quality per base > 15), and drop reads shorter than 20 bases long. Then, bacterial reads were assigned at the family level using Kraken [83]. Relative abundance of bacterial families were determined using Bracken [84]. We removed any low abundance bacterial family that was assigned fewer than 100 reads as potential contaminants.

Bacterial data was analyzed using the phyloseq package [85]. To assess if bacterial communities were fully sampled, rarefaction was performed (step size =1000) using ggrare [86]. Rarefaction curves indicate communities were fully sampled in all experiments (Supp. Figs 1–10). Beta-diversity to test differences in community composition between control and evolved populations was performed using PERMANOVA on Jaccard similarity. Bacterial alpha-diversity was calculated using the Shannon diversity index. For each experiment, each line was subsampled with replacement to the minimum number of reads in the experiment. Diversity was calculated on this rarefied library. The subsampling was performed 100 times to minimize stochasticity and artificial inflation of diversity associated with rarefaction [87]. Diversity was then averaged across the 100 subsampling efforts and compared between control and evolved lines. We determined significance using Welch’s t-test. We then tested whether two factors were sufficient to explain variation in microbial diversity between control and evolved lines: duration of selection (i.e., the number of fly generations) and *Wolbachia* infection.

First, the duration of selection ranged from 5-605 generations. We reasoned that selection response in the microbiome might be influenced by length of selection (the longer the selection, the more divergent the microbiome between control and evolved lines). To test if the duration of selection was correlated with changes in microbial diversity, we first calculated the average microbial diversity for the control lines. We then subtracted the diversity of each evolved line from the averaged control diversity to calculate change in diversity. Because we had positive and negative changes in diversity, we used the absolute difference. We performed a linear regression between change in diversity and the log10 duration of selection, modeled as Y = a + b + e, where Y= change in diversity, a = log10 duration of selection, b = random effect of experiment, and e = residual error. Lme4 was used to perform the regression in R [88].

Given that *Wolbachia* reads frequently make up the majority of the microbial reads (Supp. Table 1), we examined if *Wolbachia* relative abundance differed between control and evolved populations. We focused on only the four studies with *Wolbachia* relative abundance >2%. Statistical significance was assessed with a Kruskal-Wallis test on Wolbachia relative abundance. Second, we tested whether or not excluding *Wolbachia* reads influenced bacterial diversity. *Wolbachia* is a facultative, intracellular bacteria transmitted exclusively from mother to offspring. *Wolbachia* has no known environmental reservoir and cannot exist outside of the host. These intracellular, maternally transmitted bacteria have the same evolutionary trajectory as the host genome. The shared transmission mode with the host genome may change the response to selection compared to the other portions of the microbiome that are acquired from the environment [5, 6]. Furthermore, recent studies suggest that *Wolbachia* may interact with other bacteria in the microbiome [55, 56]. Finally, many studies in *D. melanogaster* exclude *Wolbachia* reads when analyzing microbial communities [24, 57–60], leaving an open question as to if *Wolbachia* is considered part of the *Drosophila* microbiome. To test the effect of *Wolbachia* on our inference, we removed the Anaplasmataceae family (all of which are *Wolbachia* in flies) reads from the communities, and then recalculated Shannon diversity through the same procedure as above. Libraries were rarified to the new minimum size following *Wolbachia* removal. We compared if bacterial communities without *Wolbachia* reads significantly differed between control and evolved lines using a t-test.

## List of abbreviations

E&R: Evolve and Resequence
IIS: insulin/insulin-like signalling

## Declarations

### Ethics approval and consent to participate

Not applicable.

### Consent for publication

Not applicable.

### Availability of data and materials

The raw. fastq data are available in the NCBI Sequence Read Archive, and accession numbers can be found in Supp. Table 2. The code used to analyze data can be found https://github.com/lphenry/extgeno_e-r.

### Authors’ contributions

LPH and JFA designed research. LPH collected, analyzed data and wrote the manuscript with JFA contributing to revisions. All authors read and approved the manuscript.

## Acknowledgements

We thank members of the Ayroles lab for helpful feedback.

## Supplementary information

Supplementary information accompanies this manuscript.

Additional file 1: Supp. Table 1: Relative abundance of Wolbachia in each of the 10 E&R experiments.

Additional file 2: Supp. Table 2: List of accession numbers for raw genomic data from the 10 E&R experiments.

Additional file 3: Supp. Figures 1–10: Distribution of sequencing depth, rarefaction curves, and relative abundance of bacteria (family-level) for each of the 10 E&R experiments.

Additional file 4: Supp. Figure 11: Estimates of random effects (i.e., E&R experiment) from linear model to test the correlation between microbial change and duration of selection.

**Supp. Table 1:** Relative abundance of Wolbachia in each experiment. For each experiment, the minimum, max, and average percentage of Anaplasmataceae reads is presented for both control (C) and evolved (E) lines, as well as total average for the experiment.

| Study | C-Wolb<br>avg | C-Wolb<br>min | C-Wolb<br>max | E-Wolb<br>avg | E-Wolb<br>min | E-Wolb<br>max | Average |
| --- | --- | --- | --- | --- | --- | --- | --- |
| Accelerated development | 0.02 | 0.02 | 0.03 | 1.32 | 1.24 | 1.41 | 0.67 |
| Delayed reproduction | 0.00 | 0.00 | 0.00 | 0.00 | 0.00 | 0.00 | 0.00 |
| Increased lifespan | 0.00 | 0.00 | 0.00 | 0.00 | 0.00 | 0.00 | 0.00 |
| Egg size | 0.03 | 0.03 | 0.05 | 0.06 | 0.03 | 0.12 | 0.05 |
| Desiccation resistance | 76.86 | 70.21 | 81.53 | 88.86 | 88.43 | 89.72 | 82.86 |
| Fluctuating temps | 62.51 | 48.69 | 75.35 | 72.22 | 68.45 | 75.15 | 65.42 |
| Salt + cadmium resist | 0.01 | 0.01 | 0.01 | 0.01 | 0.01 | 0.03 | 0.01 |
| Starvation resistance | 56.21 | 53.03 | 58.22 | 75.48 | 72.30 | 78.37 | 65.85 |
| Parasitoid resistance | 0.00 | 0.00 | 0.00 | 0.00 | 0.00 | 0.00 | 0.00 |
| Viral resistance | 96.39 | 95.20 | 97.31 | 97.72 | 97.57 | 98.09 | 96.84 |

**Supp. Table 2:**
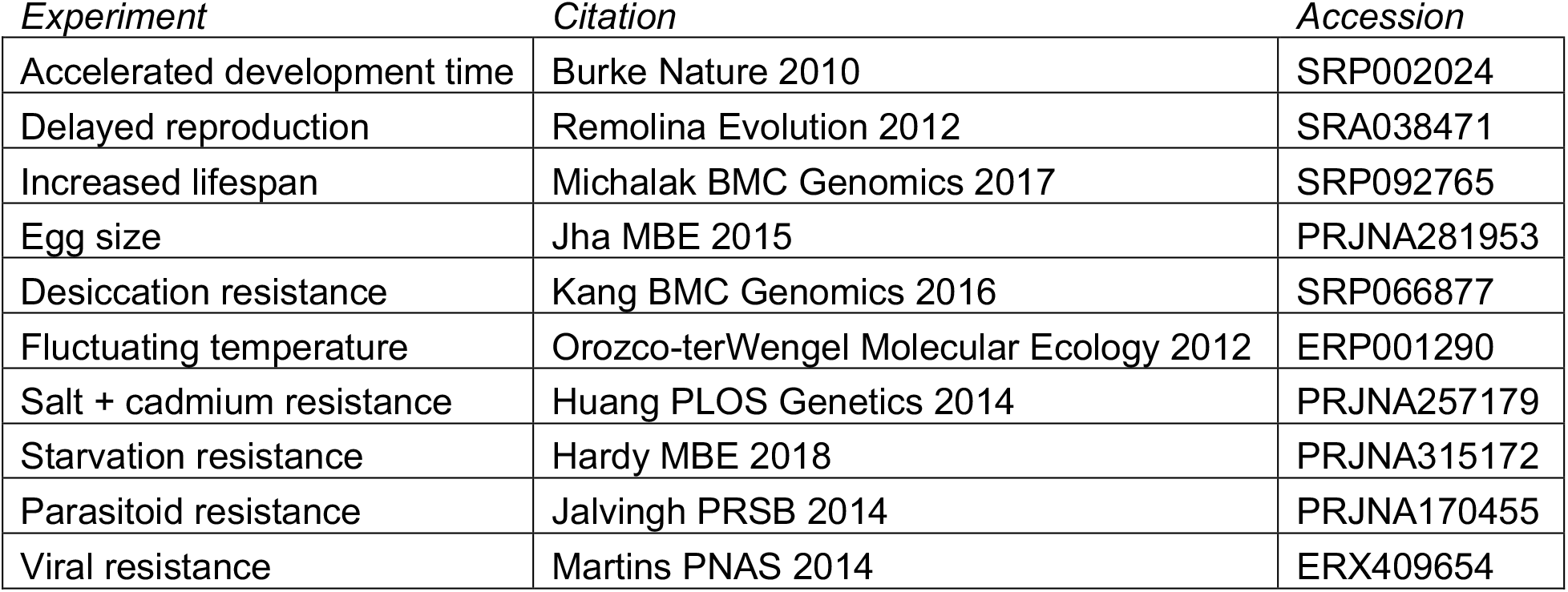
Accession numbers for E&R experiments

**Supp. Fig. 1:**
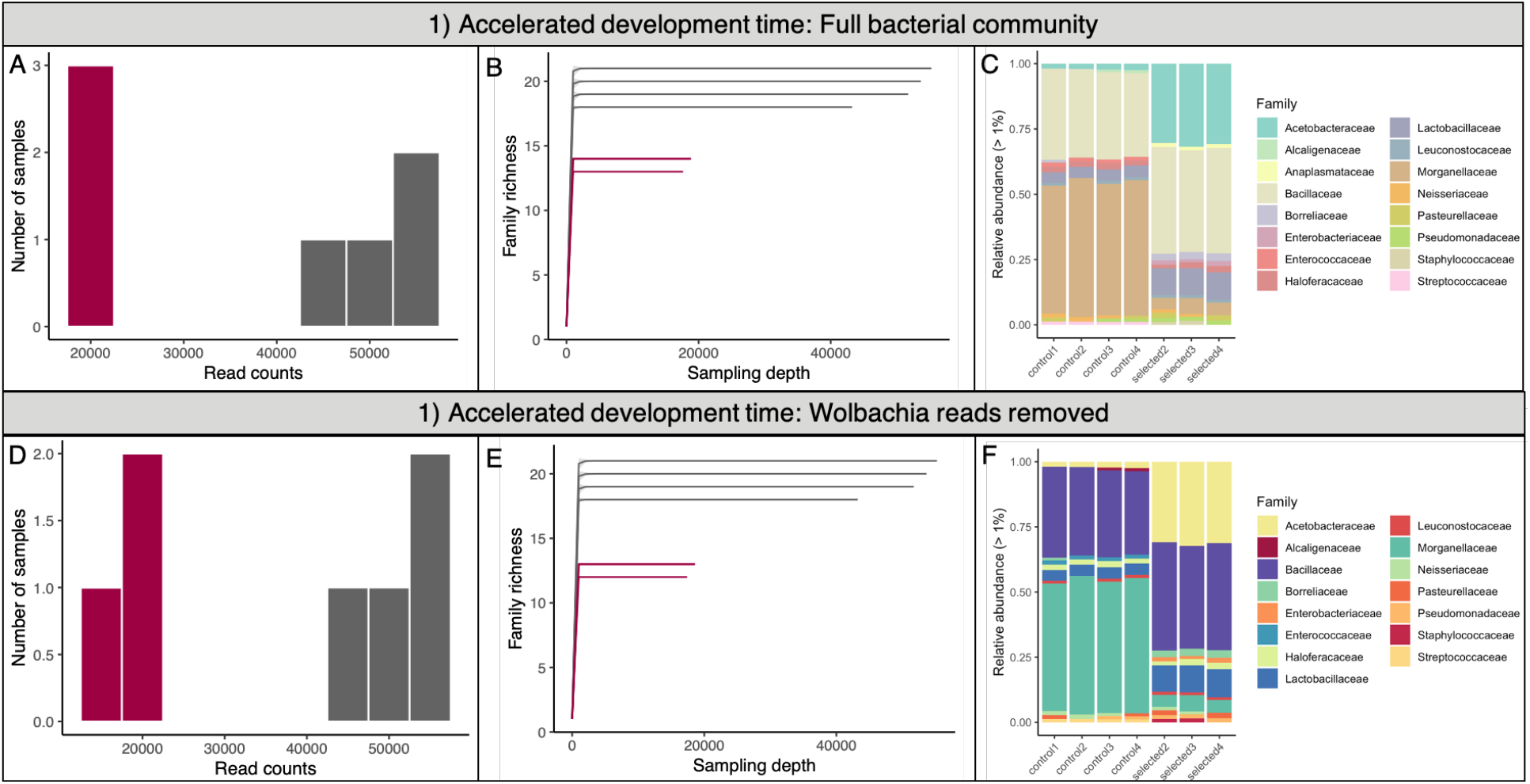
Accelerated development time. The control population is in grey, while dark red denotes the evolved populations. A) Histogram of sequencing depth for the seven pools in this study. B) Rarefaction curves suggest that bacterial communities were fully sampled, even though sequencing depth was lower for the evolved populations. C) Relative abundance of each pool shows bacterial taxa with each color. D-F) Histogram of sequencing depth, rarefaction, and relative abundance of bacterial families following removal of *Wolbachia* reads.

**Supp. Fig. 2:**
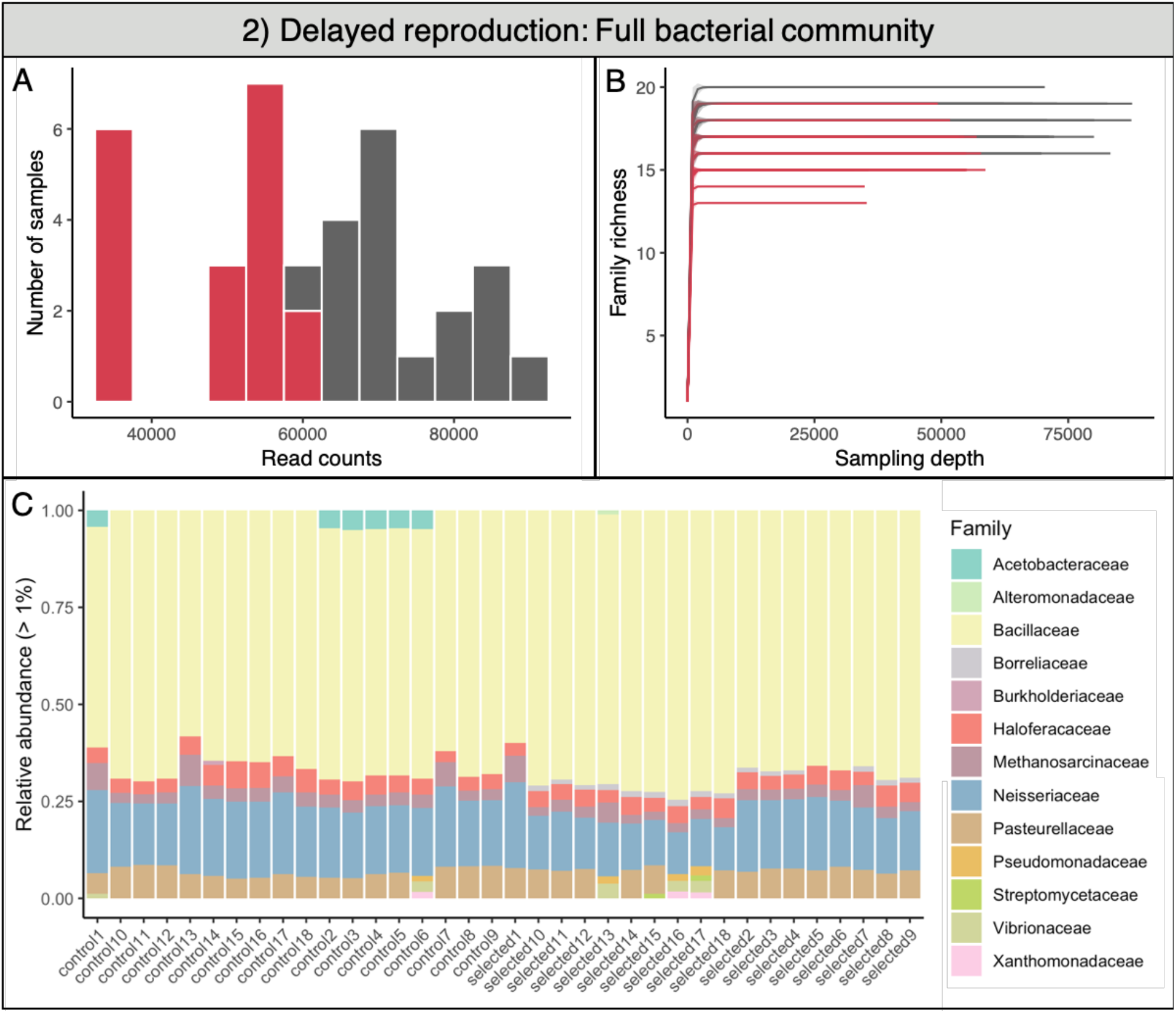
Delayed reproduction. The control population is in grey, while red denotes the evolved populations. A) Histogram of sequencing depth for the 36 pools in this study. B) Rarefaction curves suggest that bacterial communities were fully sampled, even though sequencing depth tended to be lower for the evolved populations. C) Relative abundance of each pool shows bacterial taxa with each color. *Wolbachia* was not present in this study.

**Supp. Fig. 3:**
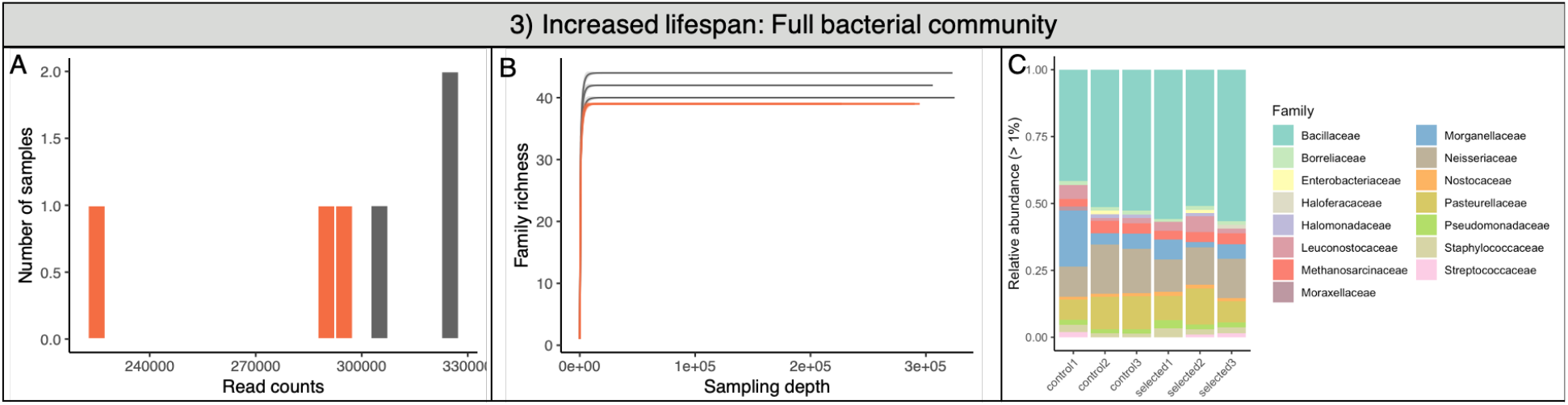
Increased lifespan. The control population is in grey, while dark orange denotes the evolved populations. A) Histogram of sequencing depth for the six pools in this study. B) Rarefaction curves suggest that bacterial communities were fully sampled, even though sequencing depth was lower for the evolved populations. C) Relative abundance of each pool shows bacterial taxa with each color. *Wolbachia* was not present in this study.

**Supp. Fig. 4:**
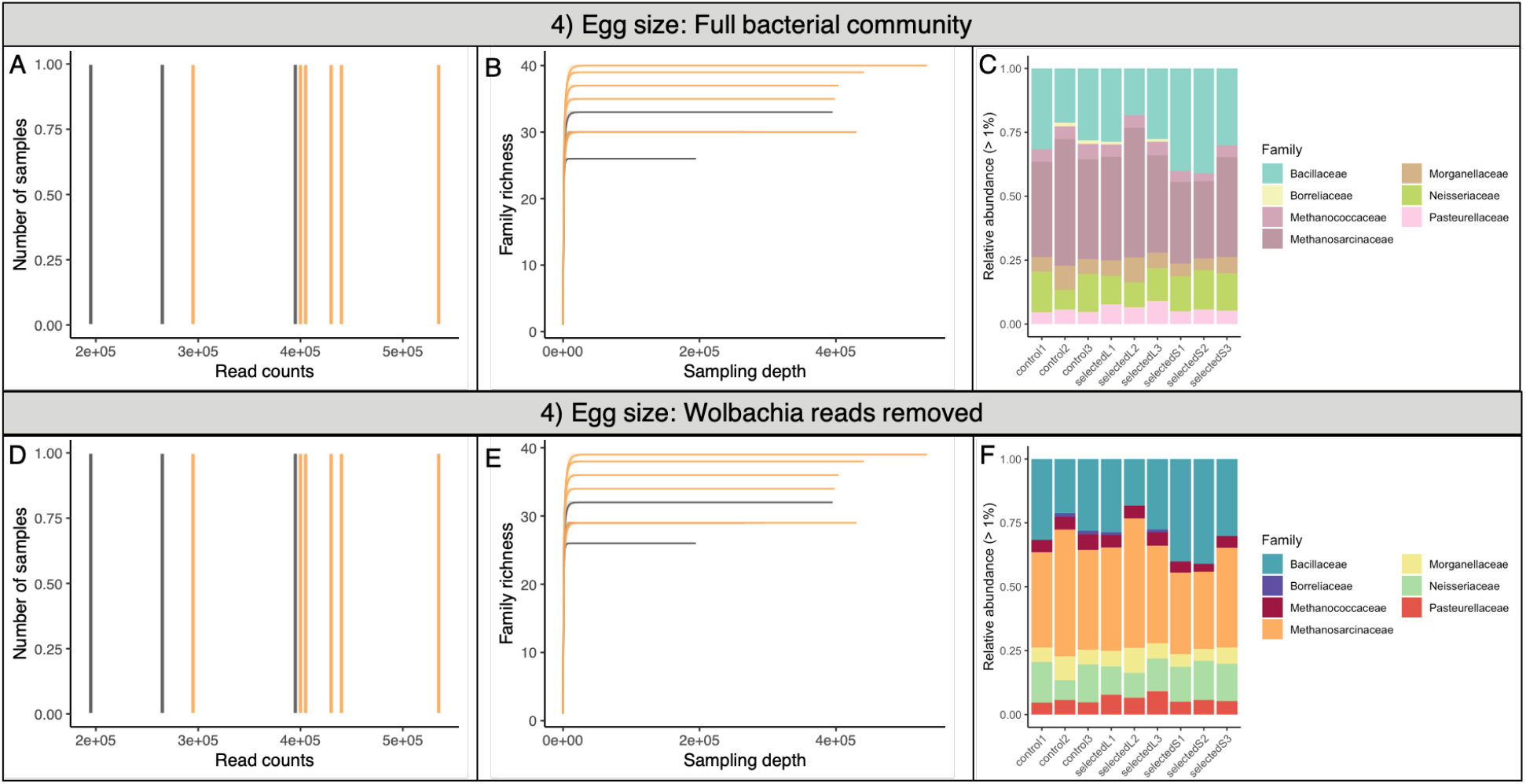
Egg size. The control population is in grey, while orange denotes the evolved populations. A) Histogram of sequencing depth for the nine pools in this study. B) Rarefaction curves suggest that bacterial communities were fully sampled, even though sequencing depth was tended to be lower for the control populations. C) Relative abundance of each pool shows bacterial taxa with each color. D-F) Histogram of sequencing depth, rarefaction, and relative abundance of bacterial families following removal of *Wolbachia* reads.

**Supp. Fig. 5:**
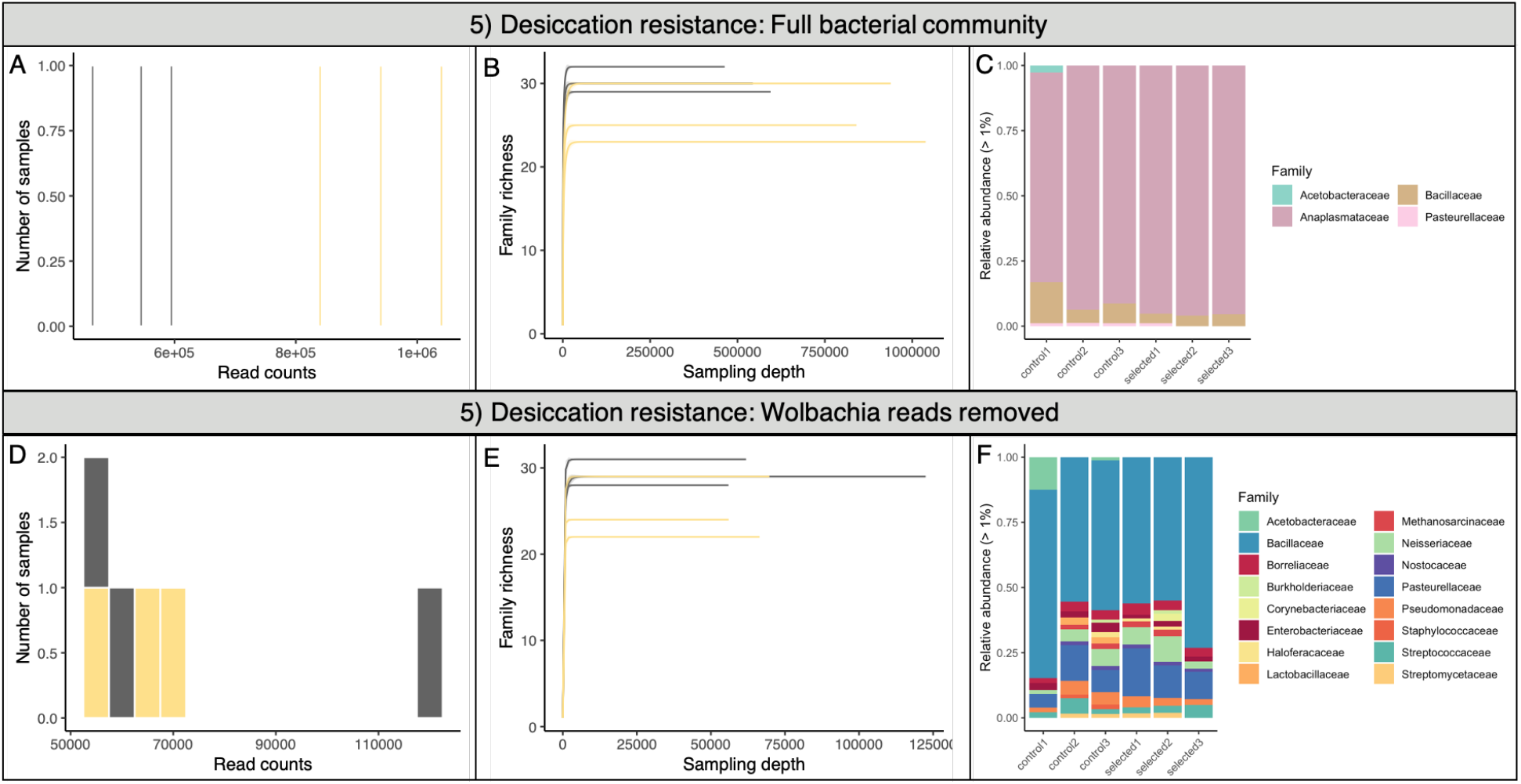
Desiccation resistance. The control population is in grey, while dark yellow denotes the evolved populations. A) Histogram of sequencing depth for the six pools in this study. B) Rarefaction curves suggest that bacterial communities were fully sampled. C) Relative abundance of each pool shows bacterial taxa with each color. D-F) Histogram of sequencing depth, rarefaction, and relative abundance of bacterial families following removal of *Wolbachia* reads.

**Supp. Fig. 6:**
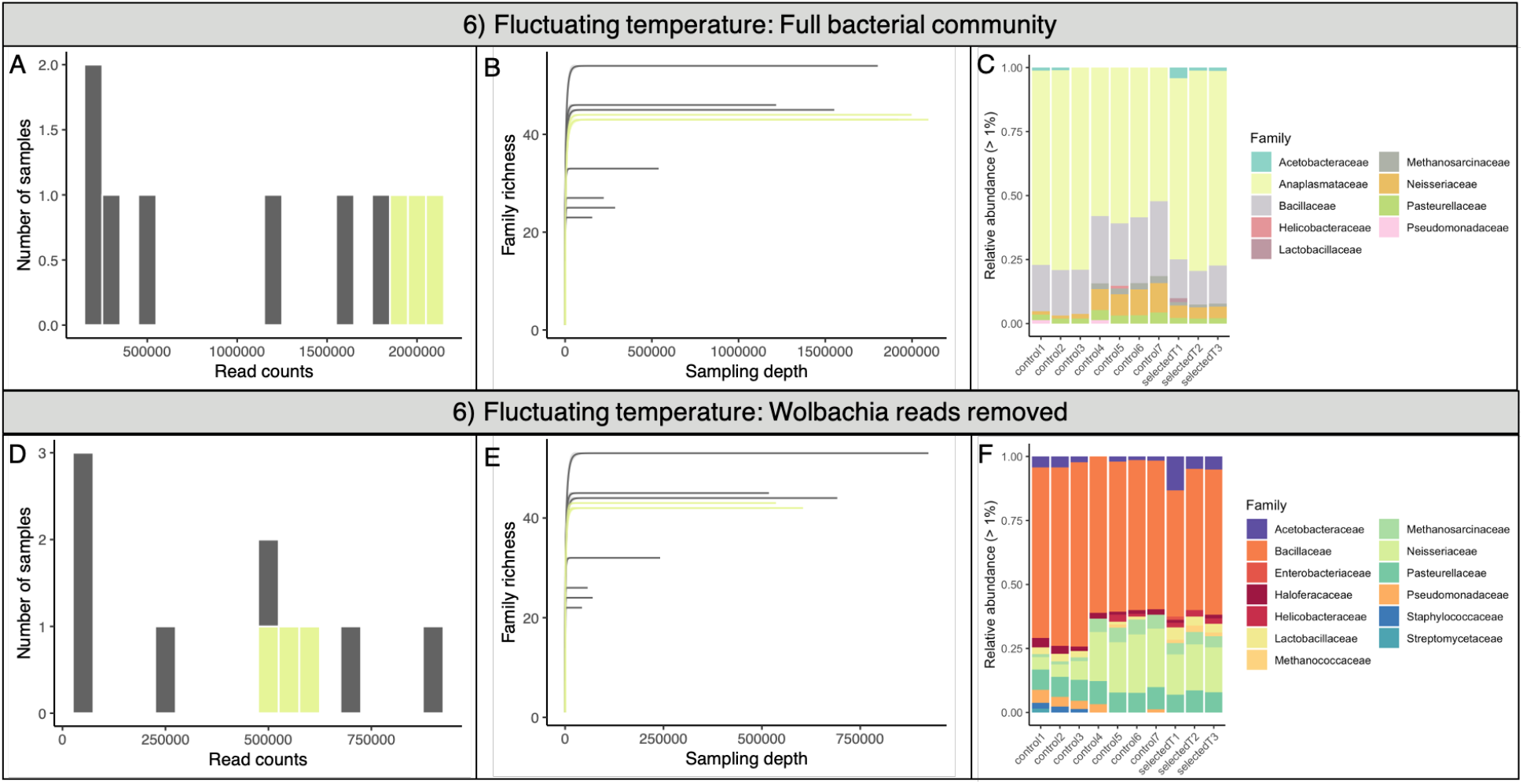
Fluctuating temperature. The control population is in grey, while yellow denotes the evolved populations. A) Histogram of sequencing depth for the 10 pools in this study. B) Rarefaction curves suggest that bacterial communities were fully sampled, even though sequencing depth was lower for the control populations. C) Relative abundance of each pool shows bacterial taxa with each color. D-F) Histogram of sequencing depth, rarefaction, and relative abundance of bacterial families following removal of *Wolbachia* reads.

**Supp. Fig. 7:**
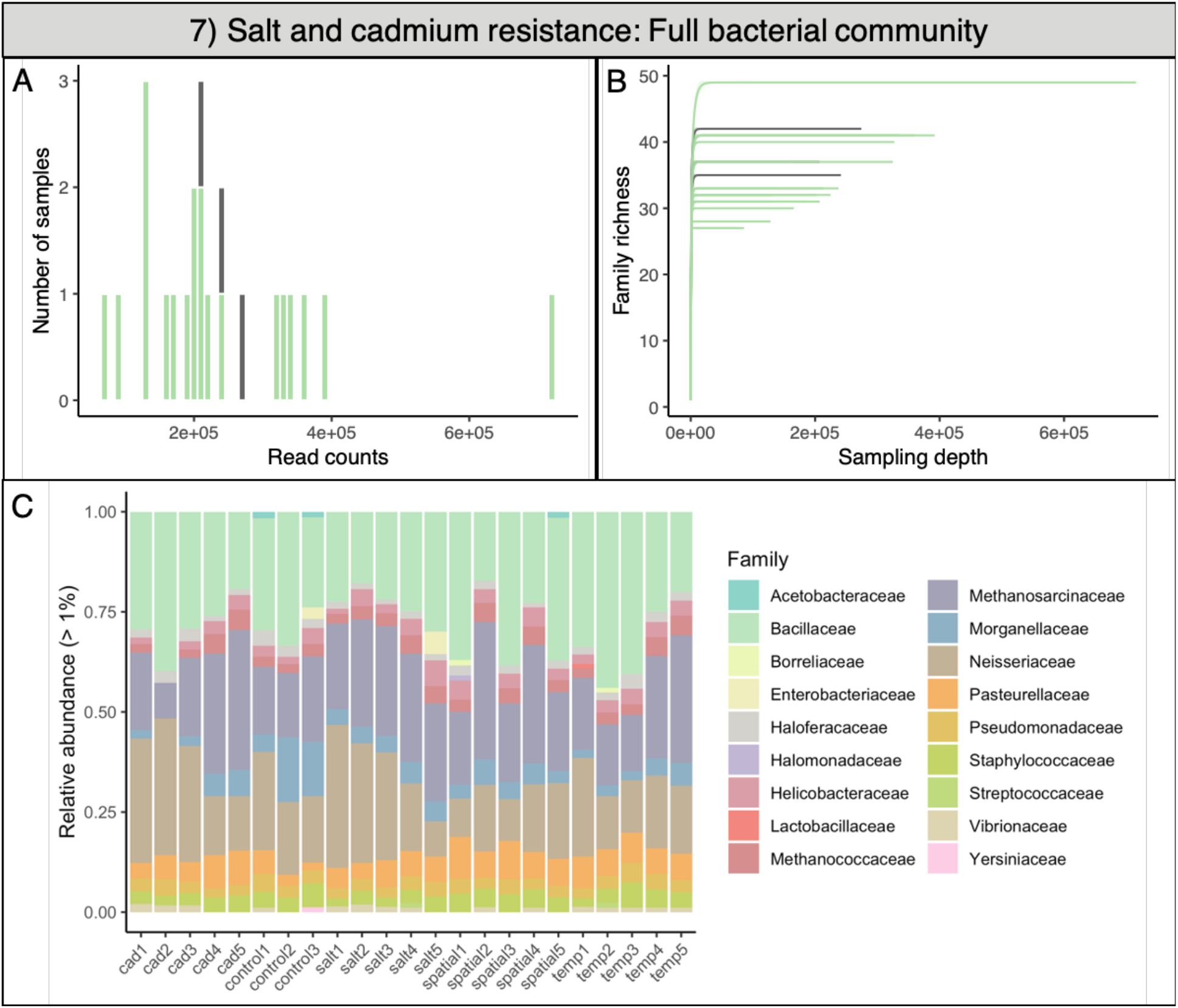
Salt and cadmium resistance. The control population is in grey, while green denotes the evolved populations. A) Histogram of sequencing depth for the 23 pools in this study. B) Rarefaction curves suggest that bacterial communities were fully sampled, even though sequencing depth was lower for the evolved populations. C) Relative abundance of each pool shows bacterial taxa with each color. While *Wolbachia* was present in this study, it was below 1% relative abundance.

**Supp. Fig. 8:**
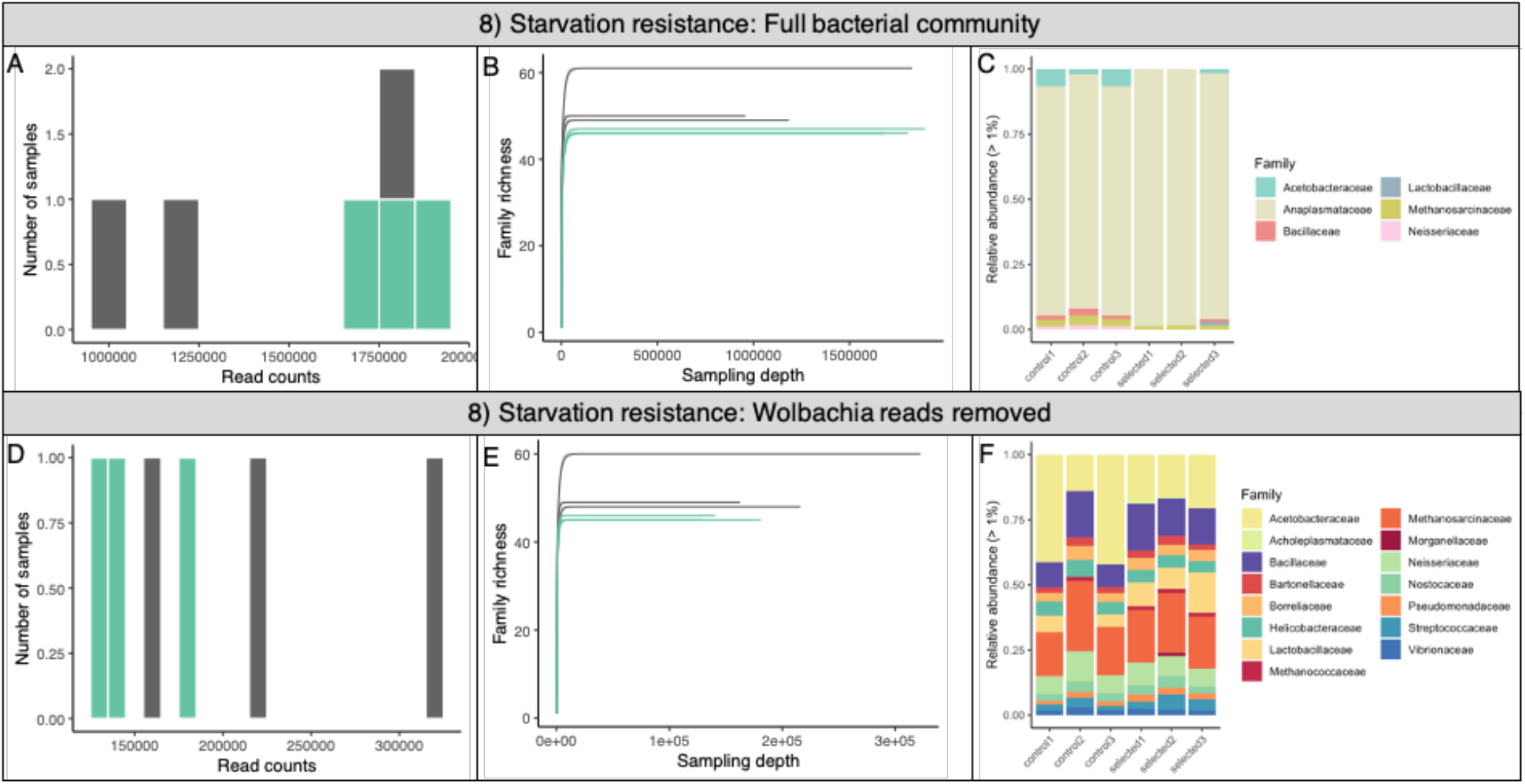
Starvation resistance. The control population is in grey, while teal denotes the evolved populations. A) Histogram of sequencing depth for the six pools in this study. B) Rarefaction curves suggest that bacterial communities were fully sampled. C) Relative abundance of each pool shows bacterial taxa with each color. D-F) Histogram of sequencing depth, rarefaction, and relative abundance of bacterial families following removal of *Wolbachia* reads.

**Supp. Fig. 9:**
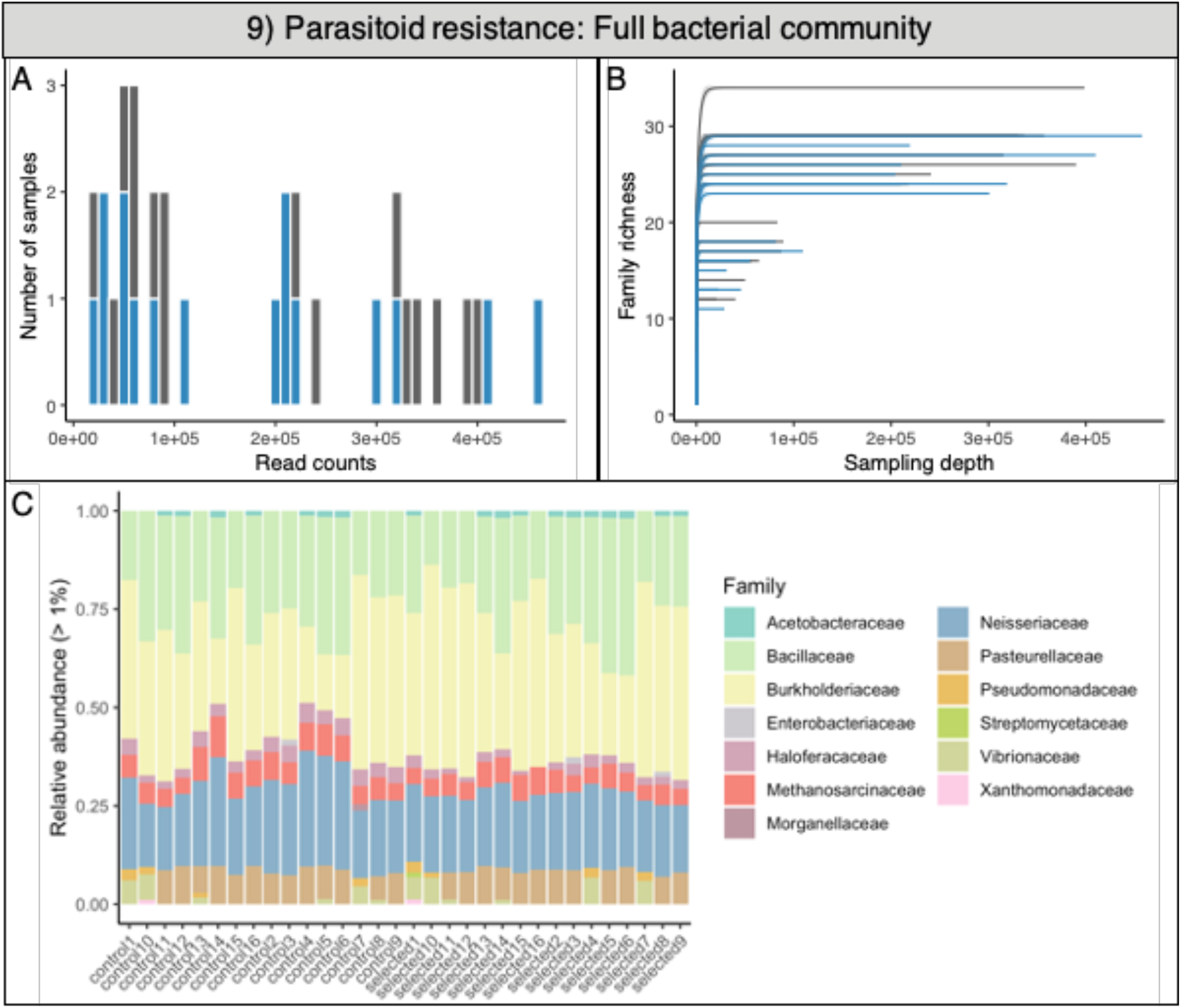
Parasitoid resistance. The control population is in grey, while blue denotes the evolved populations. A) Histogram of sequencing depth for the 24 pools in this study. B) Rarefaction curves suggest that bacterial communities were fully sampled. C) Relative abundance of each pool shows bacterial taxa with each color. *Wolbachia* was not present in this study.

**Supp. Fig. 10:**
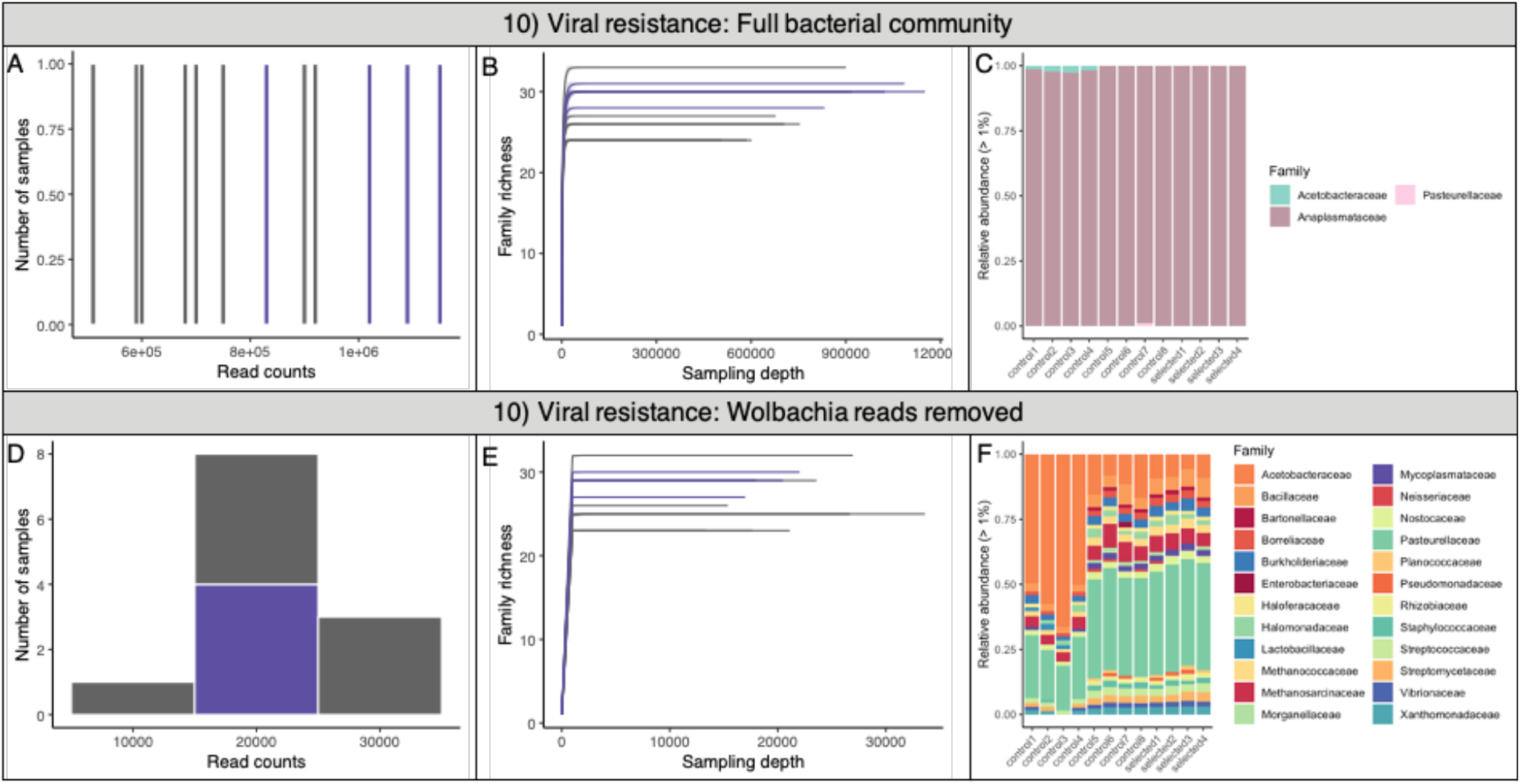
Viral resistance. The control population is in grey, while purple denotes the evolved populations. A) Histogram of sequencing depth for the 12 pools in this study. B) Rarefaction curves suggest that bacterial communities were fully sampled. C) Relative abundance of each pool shows bacterial taxa with each color. D-F) Histogram of sequencing depth, rarefaction, and relative abundance of bacterial families following removal of *Wolbachia* reads.

**Supp. Fig. 11:**
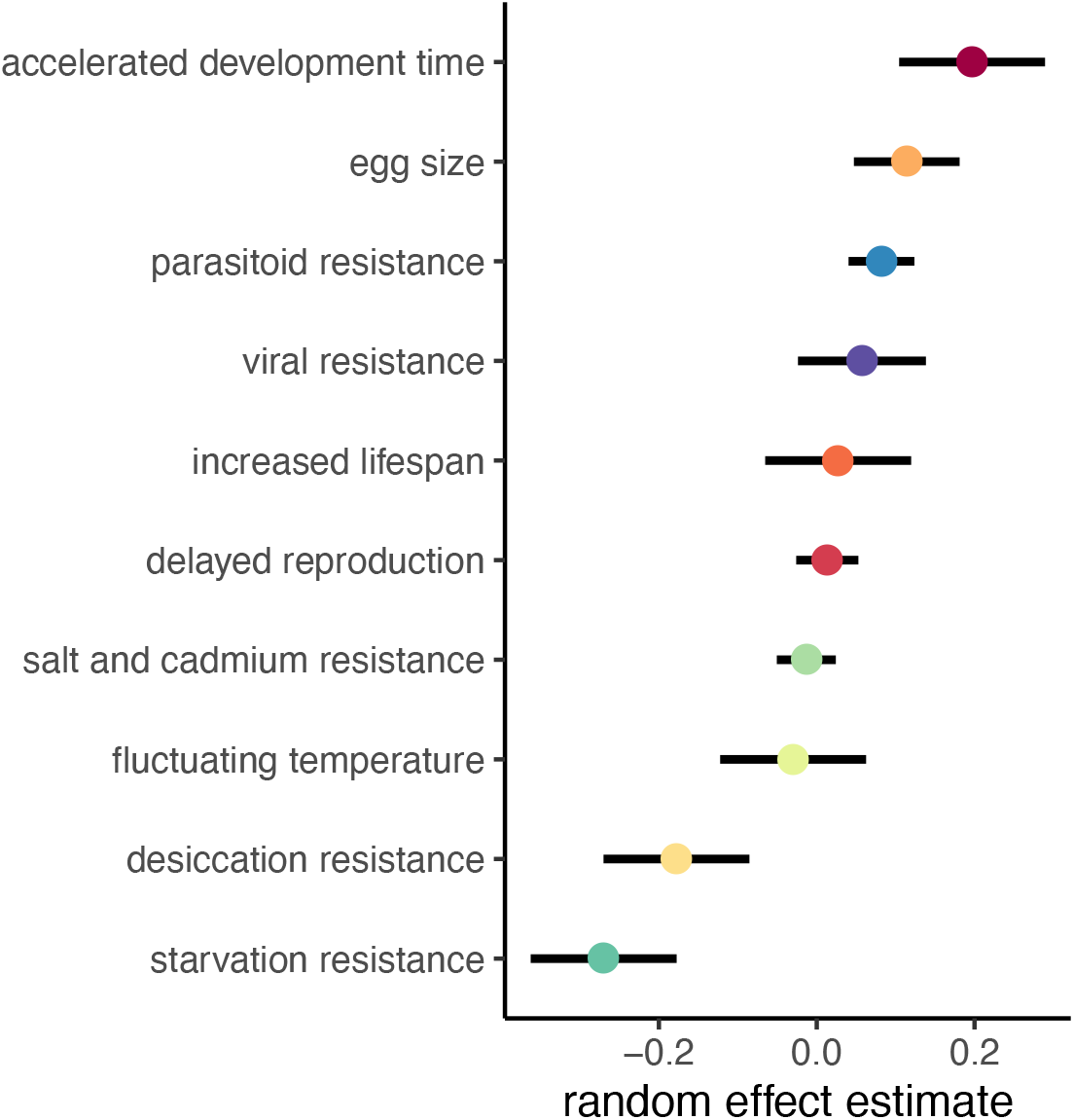
Estimates of random effects for each experiment. Points show conditional means for the random effects and lines show +/− 2 conditional SD.

## References

1. Turnbaugh PJ, Ley RE, Hamady M, Fraser-Liggett CM, Knight R, Gordon JI. The human microbiome project. Nature. 2007;449:804–10.

2. Friesen ML, Porter SS, Stark SC, von Wettberg EJ, Sachs JL, Martinez-Romero E. Microbially Mediated Plant Functional Traits. Annu Rev Ecol Evol Syst. 2011;42:23–46.

3. McFall-Ngai M, Hadfield MG, Bosch TC, Carey HV, Domazet-Loso T, Douglas AE, et al. Animals in a bacterial world, a new imperative for the life sciences. Proceedings of the National Academy of Sciences. 2013;110:3229–36.

4. Moran NA, Sloan DB. The hologenome concept: Helpful or hollow? PLoS Biol. 2015;13:e1002311.

5. Koskella B, Hall LJ, Metcalf CJE. The microbiome beyond the horizon of ecological and evolutionary theory. Nat Ecol Evol. 2017;1:1606–15.

6. Henry LP, Bruijning M, Forsberg SKG, Ayroles JF. Can the microbiome influence host evolutionary trajectories? bioRxiv. 2019;:700237.

7. Ferreiro A, Crook N, Gasparrini AJ, Dantas G. Multiscale Evolutionary Dynamics of Host-Associated Microbiomes. Cell. 2018;172:1216–27.

8. Hurst GDD. Extended genomes: symbiosis and evolution. Interface Focus. 2017;7:20170001.

9. Carthey AJR, Gillings MR, Blumstein DT. The Extended Genotype: Microbially Mediated Olfactory Communication. Trends Ecol Evol. 2018;33:885–94.

10. Mueller UG, Sachs JL. Engineering Microbiomes to Improve Plant and Animal Health. Trends Microbiol. 2015;23:606–17.

11. Hoang KL, Morran LT, Gerardo NM. Experimental Evolution as an Underutilized Tool for Studying Beneficial Animal–Microbe Interactions. Front Microbiol. 2016;07:1444.

12. Kofler R, Schlötterer C. A guide for the design of evolve and resequencing studies. Mol Biol Evol. 2014;31:474–83.

13. Long A, Liti G, Luptak A, Tenaillon O. Elucidating the molecular architecture of adaptation via evolve and resequence experiments. Nat Rev Genet. 2015;16:567–82.

14. Schlötterer C, Kofler R, Versace E, Tobler R, Franssen SU. Combining experimental evolution with next-generation sequencing: a powerful tool to study adaptation from standing genetic variation. Heredity. 2015;114:431–40.

15. Starr DJ, Cline TW. A host–parasite interaction rescues Drosophila oogenesis defects. Nature. 2002;418:76–9.

16. Clark ME, Anderson CL, Cande J, Karr TL. Widespread prevalence of Wolbachia in laboratory stocks and the implications for Drosophila research. Genetics. 2005;170:1667–75.

17. Ikeya T, Broughton S, Alic N, Grandison R, Partridge L. The endosymbiont Wolbachia increases insulin/IGF-like signalling in Drosophila. Proceedings of the Royal Society of London B: Biological Sciences. 2009;276:3799–807.

18. Tsuchida T, Koga R, Horikawa M, Tsunoda T, Maoka T, Matsumoto S, et al. Symbiotic bacterium modifies aphid body color. Science. 2010;330:1102–4.

19. Difford GF, Plichta DR, Løvendahl P, Lassen J, Noel SJ, Højberg O, et al. Host genetics and the rumen microbiome jointly associate with methane emissions in dairy cows. PLoS Genet. 2018;14:e1007580.

20. Buitenhuis B, Lassen J, Noel SJ, Plichta DR, Sørensen P, Difford GF, et al. Impact of the rumen microbiome on milk fatty acid composition of Holstein cattle. Genet Sel Evol. 2019;51:23.

21. Camarinha-Silva A, Maushammer M, Wellmann R, Vital M, Preuss S, Bennewitz J. Host Genome Influence on Gut Microbial Composition and Microbial Prediction of Complex Traits in Pigs. Genetics. 2017;206:1637–44.

22. Broderick NA, Lemaitre B. Gut-associated microbes of Drosophila melanogaster. Gut Microbes. 2012;3:307–21.

23. Douglas AE. The Drosophila model for microbiome research. Lab Anim. 2018;47:157–64.

24. Walters AW, Hughes RC, Call TB, Walker CJ, Wilcox H, Petersen SC, et al. The microbiota influences the Drosophila melanogaster life history strategy. Mol Ecol. 2020;3:639–53.

25. Burke MK, Dunham JP, Shahrestani P, Thornton KR, Rose MR, Long AD. Genome-wide analysis of a long-term evolution experiment with Drosophila. Nature. 2010;467:587–90.

26. Remolina SC, Chang PL, Leips J, Nuzhdin SV, Hughes KA. Genomic basis of aging and life-history evolution in Drosophila melanogaster. Evolution. 2012;66:3390–403.

27. Michalak P, Kang L, Sarup PM, Schou MF, Loeschcke V. Nucleotide diversity inflation as a genome-wide response to experimental lifespan extension in Drosophila melanogaster. BMC Genomics. 2017;18:84.

28. Jha AR, Miles CM, Lippert NR, Brown CD, White KP, Kreitman M. Whole-Genome Resequencing of Experimental Populations Reveals Polygenic Basis of Egg-Size Variation in Drosophila melanogaster. Mol Biol Evol. 2015;32:2616–32.

29. Kang L, Aggarwal DD, Rashkovetsky E, Korol AB, Michalak P. Rapid genomic changes in Drosophila melanogaster adapting to desiccation stress in an experimental evolution system. BMC Genomics. 2016;17:233.

30. Orozco-terWengel P, Kapun M, Nolte V, Kofler R, Flatt T, Schlötterer C. Adaptation of Drosophila to a novel laboratory environment reveals temporally heterogeneous trajectories of selected alleles. Mol Ecol. 2012;21:4931–41.

31. Huang Y, Wright SI, Agrawal AF. Genome-wide patterns of genetic variation within and among alternative selective regimes. PLoS Genet. 2014;10:e1004527.

32. Hardy CM, Burke MK, Everett LJ, Han MV, Lantz KM, Gibbs AG. Genome-Wide Analysis of Starvation-Selected Drosophila melanogaster: A Genetic Model of Obesity. Mol Biol Evol. 2018;35:50–65.

33. Jalvingh KM, Chang PL, Nuzhdin SV, Wertheim B. Genomic changes under rapid evolution: selection for parasitoid resistance. Proceedings of the Royal Society B: Biological Sciences. 2014;281:20132303–20132303.

34. Martins NE, Faria VG, Nolte V, Schlotterer C, Teixeira L, Sucena E, et al. Host adaptation to viruses relies on few genes with different cross-resistance properties. Proceedings of the National Academy of Sciences. 2014;111:5938–43.

35. Lesperance DNA, Broderick NA. Meta-analysis of Diets Used in Drosophila Microbiome Research and Introduction of the Drosophila Dietary Composition Calculator (DDCC). G3. 2020;10:2207–11.

36. Shin SC, Kim S-H, You H, Kim B, Kim AC, Lee K-A, et al. Drosophila microbiome modulates host developmental and metabolic homeostasis via insulin signaling. Science. 2011;334:670–4.

37. Chaston JM, Newell PD, Douglas AE. Metagenome-wide association of microbial determinants of host phenotype in Drosophila melanogaster. MBio. 2014;5:e01631–14.

38. White KM, Matthews MK, Hughes RC, Sommer AJ, Griffitts JS, Newell PD, et al. A metagenome-wide association study and arrayed mutant library confirm Acetobacter lipopolysaccharide genes are necessary for association with Drosophila melanogaster. G3: Genes, Genomes, Genetics. 2018;8:1119–27.

39. Selkrig J, Mohammad F, Ng SH, Chua JY, Tumkaya T, Ho J, et al. The Drosophila microbiome has a limited influence on sleep, activity, and courtship behaviors. Sci Rep. 2018;8:10646.

40. Leftwich PT, Clarke NVE, Hutchings MI, Chapman T. Reply to Obadia et al.: Effect of methyl paraben on host–microbiota interactions in Drosophila melanogaster. Proc Natl Acad Sci U S A. 2018;115:E4549–50.

41. Obadia B, Keebaugh ES, Yamada R, Ludington WB, Ja WW. Diet influences host– microbiota associations in Drosophila. Proc Natl Acad Sci U S A. 2018;115:E4547–8.

42. Sannino DR, Dobson AJ, Edwards K, Angert ER, Buchon N. The Drosophila melanogaster Gut Microbiota Provisions Thiamine to Its Host. MBio. 2018;9. doi:10.1128/mBio.00155-18.

43. Moran NA, Ochman H, Hammer TJ. Evolutionary and Ecological Consequences of Gut Microbial Communities. Annu Rev Ecol Evol Syst. 2019;50:451–75.

44. Dobson AJ, Chaston JM, Newell PD, Donahue L, Hermann SL, Sannino DR, et al. Host genetic determinants of microbiota-dependent nutrition revealed by genome-wide analysis of Drosophila melanogaster. Nat Commun. 2015;6.

45. Newell PD, Douglas AE. Interspecies Interactions Determine the Impact of the Gut Microbiota on Nutrient Allocation in Drosophila melanogaster. Appl Environ Microbiol. 2013;80:788–96.

46. Gould AL, Zhang V, Lamberti L, Jones EW, Obadia B, Korasidis N, et al. Microbiome interactions shape host fitness. Proc Natl Acad Sci U S A. 2018;115:E11951–60.

47. Fink C, Staubach F, Kuenzel S, Baines JF, Roeder T. Noninvasive Analysis of Microbiome Dynamics in the Fruit Fly Drosophila melanogaster. Appl Environ Microbiol. 2013;79:6984–8.

48. Jehrke L, Stewart FA, Droste A, Beller M. The impact of genome variation and diet on the metabolic phenotype and microbiome composition of Drosophila melanogaster. Sci Rep. 2018;8:6215.

49. Martino ME, Joncour P, Leenay R, Gervais H, Shah M, Hughes S, et al. Bacterial Adaptation to the Host’s Diet Is a Key Evolutionary Force Shaping Drosophila-Lactobacillus Symbiosis. Cell Host Microbe. 2018;24:109–19.e6.

50. Clancy DJ, Hoffmann AA. Environmental effects on cytoplasmic incompatibility and bacterial load in Wolbachia-infected Drosophila simulans. Entomol Exp Appl. 1998;86:13–24.

51. Chrostek E, Marialva MSP, Esteves SS, Weinert LA, Martinez J, Jiggins FM, et al. Wolbachia Variants Induce Differential Protection to Viruses in Drosophila melanogaster: A Phenotypic and Phylogenomic Analysis. PLoS Genet. 2013;9:e1003896.

52. Martinez J, Ok S, Smith S, Snoeck K, Day JP, Jiggins FM. Should symbionts be nice or selfish? Antiviral effects of Wolbachia are costly but reproductive parasitism is not. PLoS Pathog. 2015;11:e1005021.

53. Kriesner P, Hoffmann AA. Rapid spread of a Wolbachia infection that does not affect host reproduction in Drosophila simulans cage populations. Evolution. 2018;72:1475–87.

54. Kaur R, Martinez J, Rota-Stabelli O, Jiggins FM, Miller WJ. Age, tissue, genotype and virus infection regulate Wolbachia levels in Drosophila. Mol Ecol. 2020;29:2063–79.

55. Simhadri RK, Fast EM, Guo R, Schultz MJ, Vaisman N, Ortiz L, et al. The Gut Commensal Microbiome of Drosophila melanogaster Is Modified by the Endosymbiont Wolbachia. mSphere. 2017;2:e00287–17.

56. Ye YH, Seleznev A, Flores HA, Woolfit M, McGraw EA. Gut microbiota in Drosophila melanogaster interacts with Wolbachia but does not contribute to Wolbachia-mediated antiviral protection. J Invertebr Pathol. 2017;143:18–25.

57. Staubach F, Baines JF, Künzel S, Bik EM, Petrov DA. Host species and environmental effects on bacterial communities associated with Drosophila in the laboratory and in the natural environment. PLoS One. 2013;8:e70749.

58. Adair KL, Wilson M, Bost A, Douglas AE. Microbial community assembly in wild populations of the fruit fly Drosophila melanogaster. ISME J. 2018;12:959–72.

59. Rudman SM, Greenblum S, Hughes RC, Rajpurohit S, Kiratli O, Lowder DB, et al. Microbiome composition shapes rapid genomic adaptation of Drosophila melanogaster. Proc Natl Acad Sci U S A. 2019;116:20025–32.

60. Wang Y, Kapun M, Waidele L, Kuenzel S, Bergland A, Staubach F. Continent-wide structure of bacterial microbiomes of European Drosophila melanogaster suggests host-control. bioRxiv. 2019;:527531. doi:10.1101/527531.

61. Harcombe W, Hoffmann AA. Wolbachia effects in Drosophila melanogaster: In search of fitness benefits. J Invertebr Pathol. 2004;87:45–50.

62. Teixeira L, Ferreira Á, Ashburner M. The bacterial symbiont Wolbachia induces resistance to RNA viral infections in Drosophila melanogaster. PLoS Biol. 2008;6:e1000002.

63. Ponton F, Wilson K, Holmes A, Raubenheimer D, Robinson KL, Simpson SJ. Macronutrients mediate the functional relationship between Drosophila and Wolbachia. Proceedings of the Royal Society B: Biological Sciences. 2015;282:20142029.

64. Fry AJ, Rand DM, Poulin R. Wolbachia interactions that determine drosophila melanogaster survival. Evolution. 2002;56:1976–81.

65. Brinker P, Fontaine MC, Beukeboom LW, Falcao Salles J. Host, Symbionts, and the Microbiome: The Missing Tripartite Interaction. Trends Microbiol. 2019;27:480–8.

66. Sommer AJ, Newell PD. Metabolic basis for mutualism between gut bacteria and its impact on the Drosophila melanogaster host. Appl Environ Microbiol. 2019. https://aem.asm.org/content/85/2/e01882-18.abstract.

67. Obadia B, Güvener ZT, Zhang V, Ceja-Navarro JA, Brodie EL, Ja WW, et al. Probabilistic Invasion Underlies Natural Gut Microbiome Stability. Curr Biol. 2017;27:1999–2006.e8.

68. Douglas AE. Contradictory Results in Microbiome Science Exemplified by Recent Drosophila Research. mBio. 2018;9:01758–18.

69. Knight R, Vrbanac A, Taylor BC, Aksenov A, Callewaert C, Debelius J, et al. Best practices for analysing microbiomes. Nat Rev Microbiol. 2018;16:410–22.

70. Garud NR, Pollard KS. Population Genetics in the Human Microbiome. Trends Genet. 2020;36:53–67.

71. Lau JA, Lennon JT. Rapid responses of soil microorganisms improve plant fitness in novel environments. Proc Natl Acad Sci U S A. 2012;109:14058–62.

72. Panke-Buisse K, Poole AC, Goodrich JK, Ley RE, Kao-Kniffin J. Selection on soil microbiomes reveals reproducible impacts on plant function. ISME J. 2015;9:980–9.

73. Lauder AP, Roche AM, Sherrill-Mix S, Bailey A, Laughlin AL, Bittinger K, et al. Comparison of placenta samples with contamination controls does not provide evidence for a distinct placenta microbiota. Microbiome. 2016;4:29.

74. Pollock J, Glendinning L, Wisedchanwet T, Watson M. The Madness of Microbiome: Attempting To Find Consensus “Best Practice” for 16S Microbiome Studies. Appl Environ Microbiol. 2018;84:e02627–17.

75. de Goffau MC, Lager S, Sovio U, Gaccioli F, Cook E, Peacock SJ, et al. Human placenta has no microbiome but can contain potential pathogens. Nature. 2019;572:329–34.

76. Chandler JA, Lang JM, Bhatnagar S, Eisen JA, Kopp A. Bacterial communities of diverse Drosophila species: ecological context of a host-microbe model system. PLoS Genet. 2011;7:e1002272.

77. Ren C, Webster P, Finkel SE, Tower J. Increased Internal and External Bacterial Load during Drosophila Aging without Life-Span Trade-Off. Cell Metab. 2007;6:144–52.

78. Koyle ML, Veloz M, Judd AM, Wong AC-N, Newell PD, Douglas AE, et al. Rearing the Fruit Fly Drosophila melanogaster Under Axenic and Gnotobiotic Conditions. J Vis Exp. 2016:e54219.

79. Erkosar B, Kolly S, van der Meer JR, Kawecki TJ. Adaptation to Chronic Nutritional Stress Leads to Reduced Dependence on Microbiota in Drosophila melanogaster. MBio. 2017;8:e01496–17.

80. Cavigliasso F, Dupuis C, Savary L, Spangenberg JE, Kawecki TJ. Experimental evolution of post-ingestive nutritional compensation in response to a nutrient-poor diet. Proceedings of the Royal Society B: Biological Sciences. 2020;287:20202684.

81. Goodrich JK, Di Rienzi SC, Poole AC, Koren O, Walters WA, Caporaso JG, et al. Conducting a microbiome study. Cell. 2014;158:250–62.

82. Bolger AM, Lohse M, Usadel B. Trimmomatic: a flexible trimmer for Illumina sequence data. Bioinformatics. 2014;30:2114–20.

83. Wood DE, Salzberg SL. Kraken: ultrafast metagenomic sequence classification using exact alignments. Genome Biol. 2014;15:R46.

84. Lu J, Breitwieser FP, Thielen P, Salzberg SL. Bracken: Estimating species abundance in metagenomics data. 2016:e104.

85. McMurdie PJ, Holmes S. phyloseq: an R package for reproducible interactive analysis and graphics of microbiome census data. PLoS One. 2013;8:e61217.

86. Kandlikar GS, Gold ZJ, Cowen MC, Meyer RS, Freise AC, Kraft NJB, et al. ranacapa: An R package and Shiny web app to explore environmental DNA data with exploratory statistics and interactive visualizations. F1000Res. 2018;7:1734.

87. McMurdie PJ, Holmes S. Waste not, want not: why rarefying microbiome data is inadmissible. PLoS Comput Biol. 2014;10:e1003531.

88. Bates D, Sarkar D, Bates MD, Matrix L. The lme4 package. R package version. 2007;2:74.

